# Antibody-based vaccine for tuberculosis: validation in horse foals challenged with the TB-related pathogen *Rhodococcus equi*

**DOI:** 10.1101/292946

**Authors:** C. Cywes-Bentley, J. N. Rocha, A. I. Bordin, M. Vinacur, S. Rehman, T.S. Zaidi, M. Meyer, S. Anthony, M. Lambert, D. R. Vlock, S. Giguère, N. D. Cohen, G. B. Pier

## Abstract

Immune correlates for protection against *Mycobacterium tuberculosi*s (Mtb) infection and other intracellular pathogens are largely undetermined. Whether there is a role for antibody-mediated immunity is controversial. *Rhodococcus equi* is an intracellular pathogen causing severe pneumonia in young horse foals, eliciting a disease with many similarities to TB including intracellular residence, formation of granulomas and induction of severe respiratory distress. No purified vaccine antigens exist for *R. equi* or Mtb infections. Both express the microbial surface polysaccharide antigen poly-*N*-acetyl glucosamine (PNAG). In a randomized, controlled, blinded challenge trial, vaccination of pregnant mares with a synthetic PNAG oligosaccharide conjugated to tetanus toxoid elicited antibody that transferred to foals via colostrum and provided nearly complete protection against *R. equi* pneumonia. Infusion of PNAG-hyperimmune plasma protected 100% of foals against *R. equi* pneumonia. Vaccination induced opsonic antibodies that killed extracellular and intracellular *R. equi* and other intracellular pathogens. Killing of intracellular organisms was dependent on antibody recognition of surface expression of PNAG on infected macrophages, complement deposition and PMN-assisted lysis of infected macrophages. Protection also correlated with PBMC release of interferon-γ in response to PNAG. Antibody-mediated opsonic killing and interferon-γ release in response to PNAG may protect against disease caused by intracellular bacterial pathogens.

## INTRODUCTION

*Mycobacterium tuberculosis* (Mtb) infection is now considered the greatest cause of serious worldwide infection and associated death from a single microbe, with a clear major need for new prophylactic and therapeutic interventions (*1*). Correlates of human cellular and humoral immunity to Mtb capable of informing vaccine development are unknown, and protection studies to date, primarily in laboratory rodents and non-human primates, have not led to an effective human vaccine (*2, 3*) outside of the limited efficacy of the Bacillus Calmette-Guerin whole-cell vaccine (*3-5*). *Rhodococcus equi* is a Gram-positive, facultative intracellular pathogen that primarily infects alveolar macrophages of foals following inhalation, resulting in a granulomatous pneumonia that is pathologically similar to that caused by Mtb in humans (*6*). Like Mtb, *R. equi* also causes extrapulmonary disorders including osseous and intra-abdominal lymphadenitis (*6-8*). The disease is of considerable importance to the equine industry (*6, 8*), and has relevance to human health both as a cause of granulomatous pneumonia (*9*), and as a platform for studying Mtb infections, due to the similar pathological and clinical findings (*6-8, 10*). Additionally, both Mtb and *R. equi* synthesize the conserved surface capsule-like polysaccharide, poly-*N*-acetyl glucosamine (PNAG), a target for the development of a broadly-protective vaccine against many pathogens (*11, 12*). These findings indicate that a study evaluating vaccination against PNAG for induction of protective immunity to *R. equi* challenge in foals could provide critical information and support for testing a PNAG vaccine or PNAG-specific monoclonal antibody against human Mtb infections.

To justify the premises underlying a *R. equi* vaccine study, multiple considerations regarding immunity to intracellular pathogens, pathogenesis of Mtb and *R. equi* infections, and properties of natural and vaccine-induced immunity to PNAG had to be taken into account. Many investigators consider effective immunologic control of these intracellular pathogens to be primarily based on cell-mediated immune (CMI) responses (*13*), a finding clearly in evidence for *R. equi*. Disease occurs almost exclusively in foals less than 6 months of age, but by ~ 9 months of age most young horses become highly resistant to this pathogen (*6, 8*). This acquired natural resistance is obviously not antibody-mediated inasmuch as the solid immunity to infection in healthy horses > 9 months of age, which obviously includes pregnant mares, is not transferred to susceptible foals via antibody in the colostrum. Colostrum is the only source of maternal antibody in foals and the offspring of other animals producing an epitheliochorial placenta. Human development of TB often correlates with defective CMI (*1, 14*), particularly in those infected with the human immunodeficiency virus (*14, 15*). Nonetheless, considerable evidence has emerged to indicate that antibodies to Mtb have the potential to be major mediators of protective immunity (*16-20*). Both Mtb and *R. equi* survive within alveolar macrophages and induce granulomas, supporting the use of *R. equi* as a relevant model for TB pathogenesis, immunity, and vaccine development.

In regards to targeting the conserved PNAG surface polysaccharide, the production of this antigen by many microbes induces natural IgG antibody in most humans and animals (*21, 22*), but natural antibody is generally ineffective at eliciting protection against infection. These antibodies do not activate the complement pathway and cannot mediate microbial killing (*21-23*). By removing most of the acetate substituents from the *N*-acetyl glucosamine sugars comprising PNAG (*24, 25*), or using synthetic oligosaccharides composed of only β-1→6-linked glucosamine conjugated to a carrier protein such as tetanus toxoid (TT), (*11, 23, 26, 27*) complement-fixing, microbial-killing, and protective antibody to PNAG can be induced. A final premise justifying evaluation of vaccine-induced immunity to *R. equi* by immunizing pregnant mares is that foals are considered to be infected soon after birth (*28*) when they are more susceptible to infection (*29*) and when their immune system is less effective in responding to vaccines, (*30-33*) which precludes active immunization of very young foals as a strategy for vaccine evaluation against *R. equi*.

Therefore, in order to ascertain if *R. equi* pneumonia could be prevented by antibody to PNAG, pregnant mares were vaccinated with the 5GlcNH_2_-TT vaccine, the transfer of functional opsonic antibodies via colostrum to foals verified, and foals were then challenged at 25-28 days of life with virulent *R. equi.* This approach protected 11 of 12 (92%) foals born to immunized mares against essentially all signs of clinical pneumonia, whereas 6 of 7 (86%) foals born to non-immune, control mares developed *R. equi* pneumonia. A follow-up study in which 2 liters (approximately 40 ml/kg) of hyperimmune plasma from adult horses immunized with the 5GlcNH_2_-TT vaccine were infused into newborn foals on day 1 of life similarly protected 5 of 5 (100%) of the recipients from *R. equi* pneumonia following challenge at 4 weeks of life whereas recipients of control plasma all developed clinical signs of pneumonia (4/4). *In vitro* correlates of immunity were further investigated, and a novel mechanism of killing of intracellular *R. equi* and other intracellular bacterial pathogens was identified, wherein intracellular bacterial growth resulted in high levels of surface-expressed PNAG intercalated into the plasma membrane of infected host cells, which served as a target for the complement-fixing antibody to PNAG, which, along with added polymorphonuclear neutrophils (PMNs), lysed the infected cell and released the intracellular organisms that were further opsonized and killed in this setting. The results establish a solid basis for evaluating vaccination or MAb therapy against PNAG for prevention or treatment of human TB and provide a clear immunologic rationale for antibody-mediated protection against intracellular pathogens.

## RESULTS

### Maternal vaccination induces serum and colostral antibody to PNAG that is orally transferred to foals

Mares were immunized twice approximately 6 and 3 weeks prior to their estimated date of parturition (based on last known breeding date) with 125 or 200 µg of the 5GIcNH_2_ vaccine conjugated to TT (AV0328 from Alopexx Vaccine, LLC) adjuvanted with 100 µl of Specol. Immunization of mares resulted in no detectable local or systemic reaction following either 1 or 2 vaccine doses except for a slightly swollen muscle 24 h after the first vaccination followed at day 2 by a small dependent edema that resolved by day 3 in a single mare. Serum samples from mares immunized in 2015 were only collected on the day of foaling, so statistical comparisons with immunized mare titers were only made between all 7 control samples collected on the day of foaling (D0 post-foaling (PF)) with 12 vaccinated samples collected pre-immunization, on day 21 prior to the booster dose, and on D0 PF (Fig. S1). When compared with IgG titers to PNAG in non-immune controls obtained on D0 PF, immunization of mares gave rise to significant (P < 0.05) increases in total IgG to PNAG as well as the equine IgG subisotypes IgG_1_, IgG_3/5_, and IgG_4/7_ in serum (Fig. S1) on day 21 after a single immunization, as well as on D0 PF, representing the serum sample obtained after the booster immunization. Similarly, total IgG and IgG subisotype titers were significantly higher in the colostrum obtained on the day of foaling from vaccinated mares compared with controls (Fig. S2). Notably, non-immunized mares had antibody titers to PNAG, representative of the natural response to this antigen commonly seen in normal animal and human sera.

Successful oral delivery of antibody to the blood of foals born to vaccinated mares (hereafter termed vaccinated foals) was shown by the significantly higher titers of serum IgG to PNAG compared with foals from control mares at ages 2, 28, and 56 days, but not 84 days (Fig. 1A). Foal serum concentrations of subisotypes IgG_1_, IgG_3/5_, and IgG_4/7_ to PNAG were significantly higher at 2, 28, and 42 days of age (after colostral transfer) in the vaccinated group compared with the control group, and subisotype IgG_1_ titers remained significantly higher through age 56 days (Figs. 1B-1D). The pattern in vaccinated foals of decreasing titers to PNAG with increasing age was consistent with the decay of maternally-transferred immunoglobulins.

**Fig 1.**
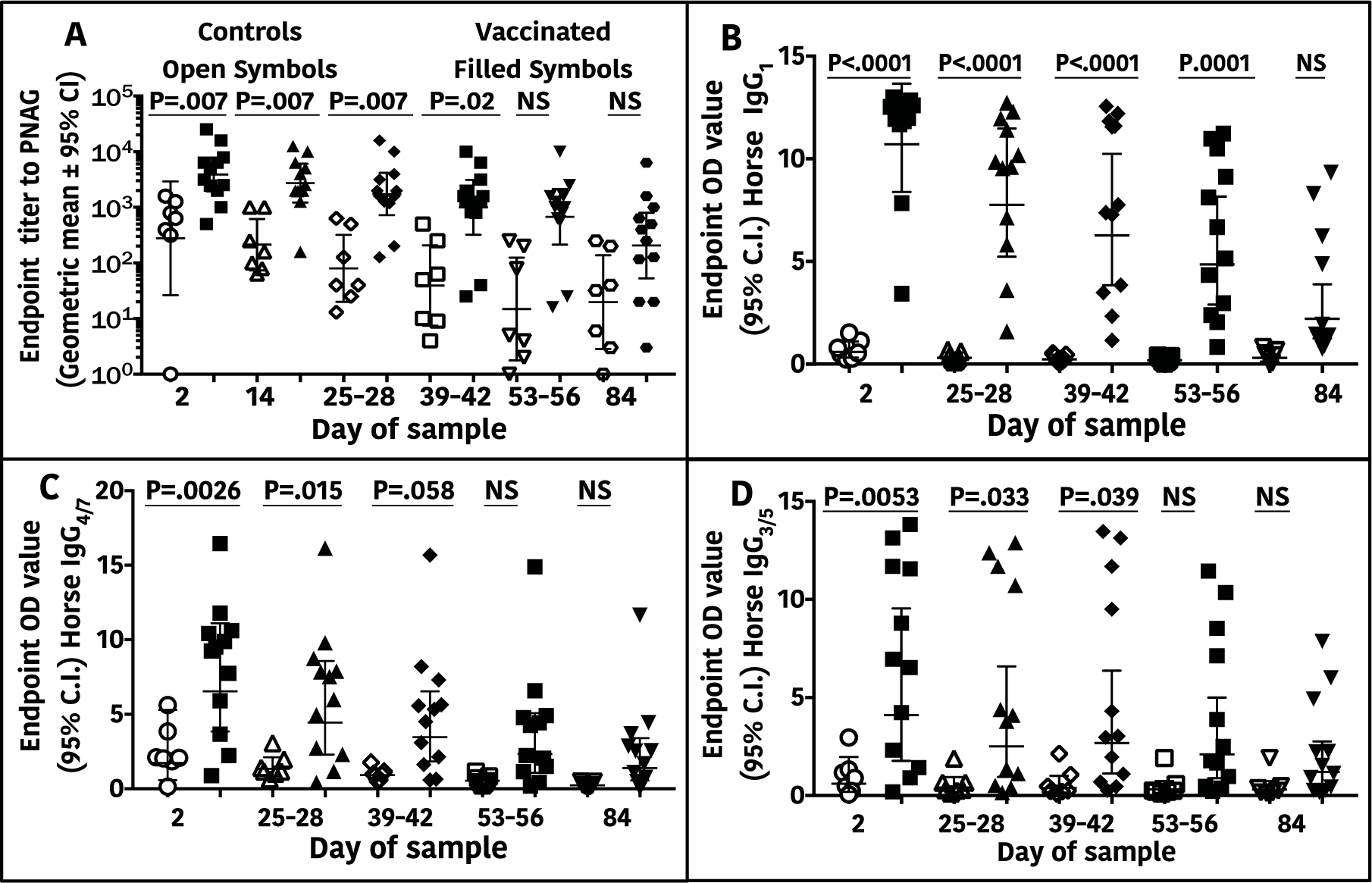
Total IgG and IgG subisotype antibody titers to PNAG in sera of horse foals. Endpoint serum titers (N=7 Non-vaccinated, 12 vaccinated) of IgG or IgG subisotypes are plotted by vaccine group as a function of age in days. (**A**) Total IgG antibody end-point titers to PNAG were significantly higher in an age-dependent matter between foals from mares that were vaccinated (filled symbols n=12) compared with titers in sera of foals from unvaccinated, control mares (open symbols n=7) through D39-42 of life. (**B-D**) Concentrations of IgG_1,_ IgG_4/7_, and IgG_3/5_ were significantly higher in foals in the vaccinated group than the unvaccinated, control group through the day indicated on the figure. Statistical comparisons made using linear mixed-effects modeling with individual foal as a random effect; NS=not significant.

### Orally obtained colostral antibody to PNAG protects foals against intrabronchial infection with *R. equi*

Protection studies were undertaken using a randomized, controlled, blinded experimental trial design. At days 25-28 of life, foals in the study were challenged with 1 × 10^6^ CFU of live *R. equi* contained in 40 ml of vehicle, with half of the challenge delivered to each lung by intrabronchial dosing with 20 ml. Foals were followed for development of clinical *R. equi* pneumonia (Table S1) for 8 weeks. The proportion of vaccinated foals that developed *R. equi* pneumonia (8%; 1/12) was significantly (P = 0.0017; Fisher’s exact test) less than that of unvaccinated control foals (86%; 6/7), representing a relative risk reduction or protected fraction of 84% (95% C.I. 42% to 97%, Koopman asymptotic score analysis (*34*)). The duration of clinical signs indicative of *R. equi* pneumonia was significantly (P ≤ 0.027, Wilcoxon rank-sum tests) longer for foals from control than vaccinated mares (Table S2). Thoracic ultrasonographic examination is the standard clinical technique for monitoring areas of pulmonary abscessation or consolidation attributed to *R. equi* infection. The severity and duration of ultrasonographic lesions were significantly greater in foals born to controls than vaccinated mares (Fig. 2). Vaccinated foals that were protected against pneumonia had less severe clinical signs and smaller and fewer ultrasonographic lesions compared with control foals. Thus, maternal vaccination against PNAG demonstrated successful protection against clinical *R. equi* pneumonia, a disease for which there is no current vaccine, using a randomized, blinded experimental challenge model.

**Fig. 2.**
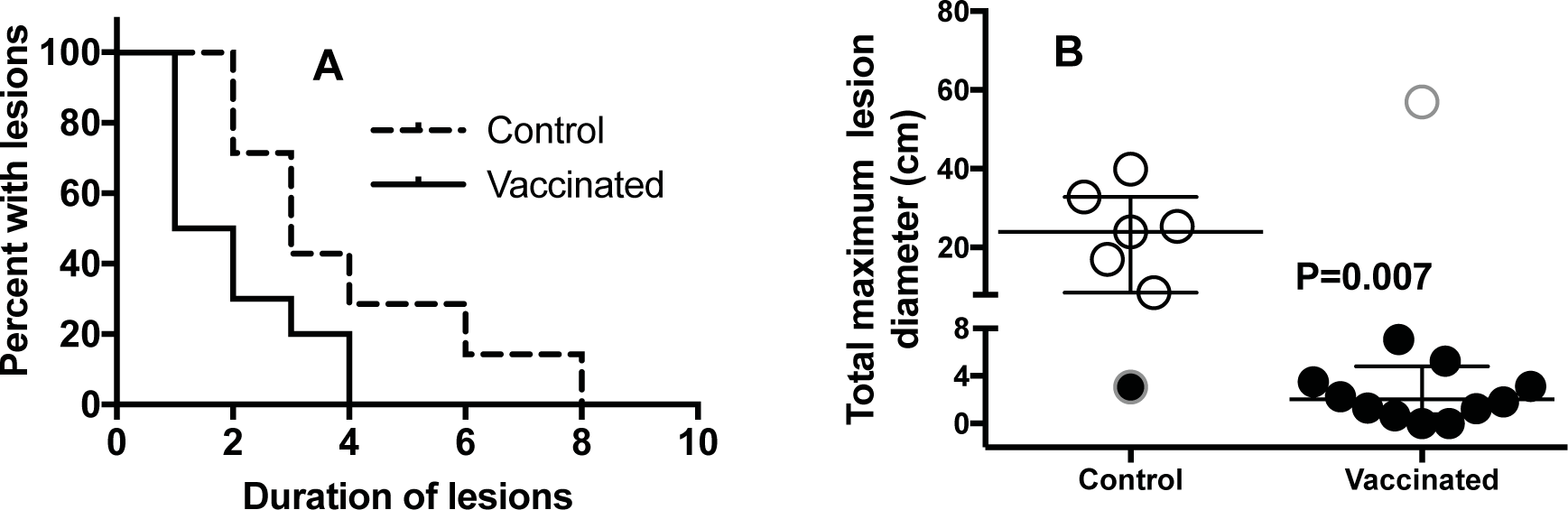
Comparison of induction and regression of ultrasonographic lesions in foals from vaccinated or unvaccinated mares following *R. equi* challenge. **(A)** Kaplan-Meier survival plot comparing duration of detectable ultrasonographic lesions as evidence of pulmonary abscessation. Duration of pulmonary lesions identified by ultrasound was significantly (P = 0.008; Log-rank test) shorter for foals of vaccinated mares (solid line) versus those of foals from control mares (hatched line). **(B)** Cumulative sum of maximum diameters of thoracic ultrasonography lesions (N=7 Unvaccinated, 12 Vaccinated). The sums of the cumulative maximum diameters were significantly (P = 0.007; Wilcoxon rank-sum test) shorter for foals from vaccinated mares (n=12) than for unvaccinated control mares (n=7). Open circles indicate foals diagnosed with pneumonia, filled circles indicate foals that did not develop pneumonia. Symbols with outer gray rings indicate the unvaccinated foal that did not get pneumonia and the vaccinated foal that did develop pneumonia.

### Passive infusion with hyperimmune plasma to PNAG protects foals against *R. equi* pneumonia

To substantiate that vaccination-mediated protection was attributable to antibody to PNAG, hyperimmune plasma was prepared from the blood of 5GlcNH_2_-TT-immunized adult horses and 2 L (approximately 40 ml/kg) infused into 5 foals at 18-24 hours of age. Four controls were transfused at the same age with 2 L of standard commercial horse plasma. Titers of control and hyperimmune plasma IgG subisotypes and IgA antibody to PNAG and OPK activity against *R. equi* (Fig. S3) documented significantly higher titers of functional antibody to PNAG in the plasma from vaccinated donors and in foals transfused with the plasma from vaccinated donors compared to foals transfused with standard plasma. After challenge with *R. equi* as described above, there was a significant reduction in clinical signs in the foals receiving PNAG-hyperimmune plasma, compared to controls, except for the duration of ultrasound lesions (Table S3). None of the 5 foals receiving PNAG-hyperimmune plasma were diagnosed with *R. equi* pneumonia, whereas 4 of 4 recipients of normal plasma had a diagnosis of clinical pneumonia for at least 1 day (P=0.0079, Fisher’s exact test; relative risk reduction or protected fraction 100%, 95% C.I. 51%-100%, Koopman asymptotic score (*34*)).

### R. equi expression of PNAG *in vitro* and *in vivo*

Using immunofluorescence microscopy, we demonstrated that 100% of 14 virulent strains of *R. equi* tested express PNAG (Fig. S4). Moreover, we found that PNAG was expressed in the lungs of foals naturally infected with *R. equi* (Fig. S5a), similar to our prior demonstration of PNAG expression in the lung of a human infected with Mtb (*11*). PNAG was detected within apparent vacuoles inside *R. equi*-infected horse macrophages *in vivo* (Fig. S5b).

### PNAG vaccine-induced opsonic antibodies mediate killing of both extracellular and intracellular *R. equi*

Testing of the functional activity of the antibodies induced in the pregnant mares and in foal sera on the day of challenge demonstrated the antibodies could fix equine complement component C1q onto the PNAG antigen (Fig. 3A). Notably, the natural antibody to PNAG in sera of non-vaccinated, control mares and their foals did not deposit C1q onto the PNAG antigen, consistent with prior findings that natural antibodies are immunologically inert in these assays (*21, 22, 35*). Sera from vaccinated foals on the day of *R. equi* infection mediated high levels of opsonic killing of extracellular *R. equi* whereas control foals with only natural maternal antibody to PNAG had no killing activity (Fig. 3B), again demonstrating the lack of functional activity of these natural antibodies to PNAG.

**Fig. 3.**
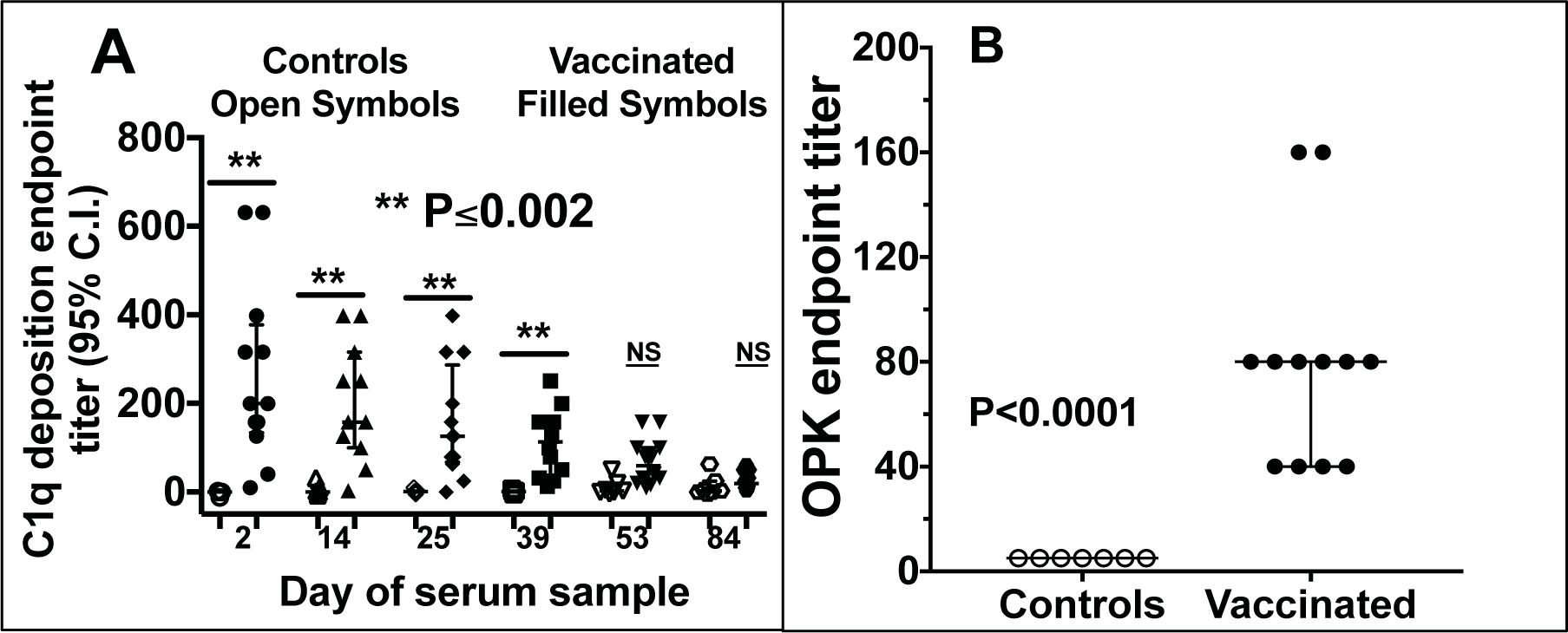
Functional activity of antibody in foal sera on day of challenge with *R. equi*. **(A)** Serum endpoint titer (N=7 unvaccinated controls, 12 vaccinated) of deposition of equine C1 onto purified PNAG. P values determined by non-parametric ANOVA and pairwise comparisons by Dunn’s procedure. NS, not significant. (**B**) Serum endpoint titer (reciprocal of serum dilution achieving killing ≥ 30% of input bacteria) for opsonic killing of *R. equi* in suspension along with horse complement and human PMN. Values indicate individual titer in foal sera on day of challenge with *R. equi*, black bars the group median and error bars the 95% C.I. (upper 95% C.I for vaccinated foals same as median). P value by Wilcoxon rank-sum test.

As some of the vaccinated foals developed small subclinical lung lesions that resolved rapidly (Table S1, Fig. 2) it appeared the bolus challenge did lead to some uptake of *R. equi* by alveolar macrophages but without development of detectable clinical signs of disease. This observation suggested that antibody to PNAG led to resolution of these lesions and prevented the emergence of clinical disease. Based on the finding that *R. equi*-infected foal lung cells expressed PNAG *in vivo* (Fig. S5), we determined if macrophages infected with *R. equi in vitro* similarly expressed PNAG, and also determined if this antigen was on the infected cell surface, intracellular, or both. We infected cultured human monocyte-derived macrophages (MDM) for 30 min with live *R. equi* then cultured them overnight in antibiotics to prevent extracellular bacterial survival. To detect PNAG on the infected cell surface we used the human IgG1 MAb to PNAG (MAb F598) conjugated to the green fluorophore Alexa Fluor 488. To detect intracellular PNAG, we next permeabilized the cells with ice-cold methanol and added either unlabeled MAb F598 or control MAb, F429 (*36*) followed by donkey anti-human IgG conjugated to Alex Fluor 555 (red color). These experiments showed there was no binding of the MAb to uninfected cells (Fig. S6A) nor binding of the control MAb to infected cells (Fig. S6B). However, we found strong expression of PNAG both on the infected MDM surface and within infected cells (Fig. S6C). Similarly, using a GFP-labeled Mtb strain (Figs. S6 D-E) and a GFP-labeled strain of *Listeria monocytogenes* (Fig. S6F) we also visualized intense surface expression of PNAG on infected human MDMs in culture, even when the bacterial burden in the infected cell was apparently low. Importantly, within infected cultures, only cells with internalized bacteria had PNAG on their surface (Fig. S6G), indicating the antigen originated from the intracellular bacteria. Thus, uninfected cells did not obtain PNAG from shed antigen or lysed infected cells. This finding is consistent with published reports of intracellular bacterial release of surface vesicles that are transported among different compartments of an infected host cell (*37*).

Next, we examined if the surface PNAG on infected cells provided the antigenic target needed by antibody to both identify infected cells and, along with complement and PMN, lyse the cells, release the intracellular microbes, and kill them by classic opsonic killing. Human MDM cultures were established *in vitro*, infected for 30 min with live *R. equi*, and then cells were washed and incubated for 24 h in the presence of 100 µg gentamicin/ml to kill extracellular bacteria and allow for intracellular bacterial growth. Then, various combinations of the human IgG_1_ MAb to PNAG or the control MAb F429 along with human complement and human PMN were added to the cultures, and viable *R. equi* determined after 90 min. While a low level of killing (≤30%) of intracellular *R. equi* was obtained with PMN and complement in the presence of the control MAb, there was a high level of killing of the intracellular *R. equi* when the full compendium of immune effectors encompassing MAb to PNAG, complement, and PMN were present (Fig. 4A). Similarly, testing of sera from vaccinated foals on the day of challenge, representing animals with a low, medium, or high titers of IgG to PNAG, showed they also mediated titer-dependent killing of intracellular *R. equi* (Fig. 4B). Measurement of the release of lactate dehydrogenase as an indicator of lysis of the macrophages showed that the combination of antibody to PNAG, complement, and PMN mediated lysis of the infected human cells (Fig. 4C), presumably releasing the intracellular bacteria for further opsonic killing.

**Fig. 4.**
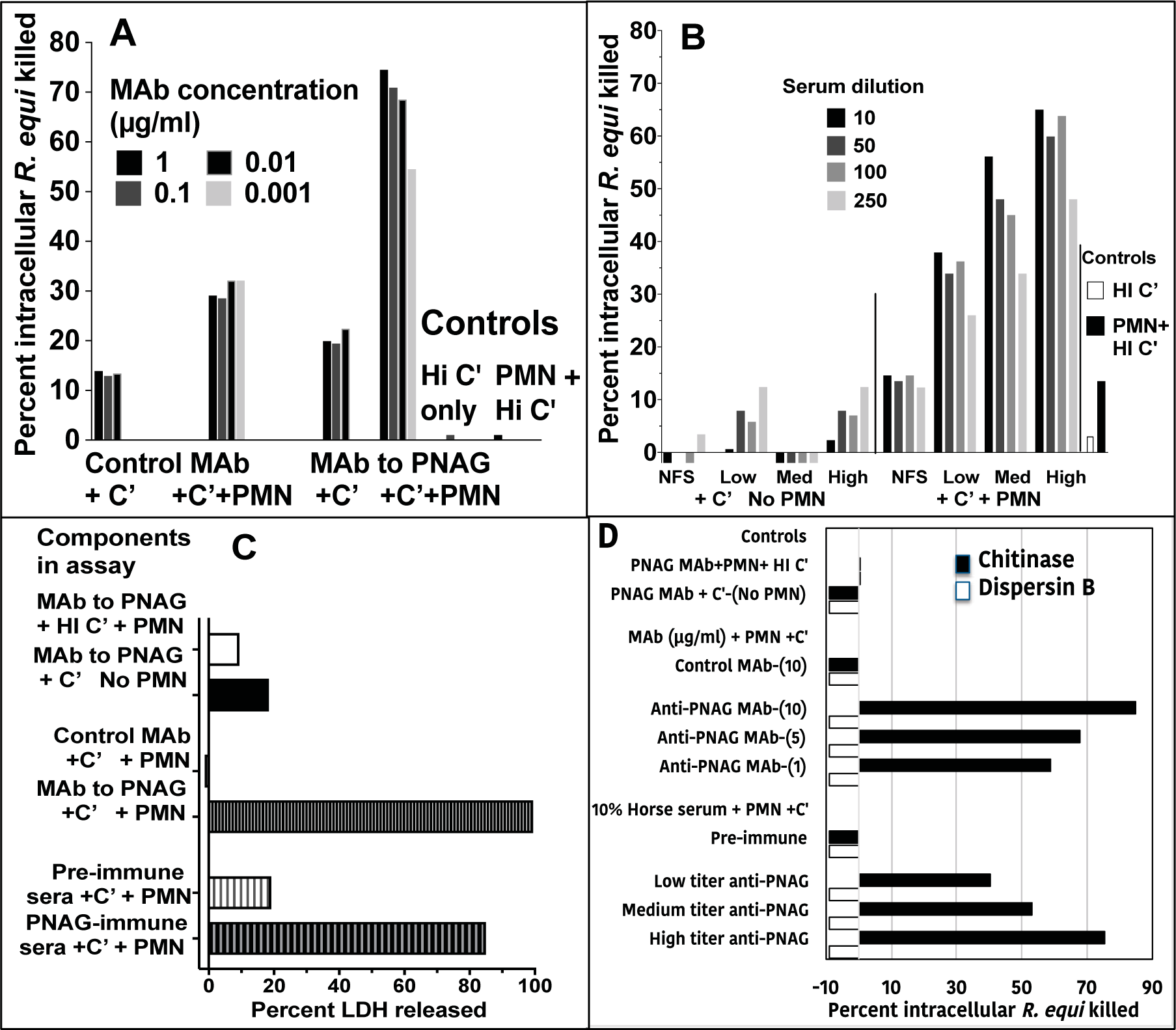
Opsonic killing of intracellular *R. equi*. (**A**). Maximal killing of intracellular *R. equi* mediated by MAb to PNAG requires both complement (C’) and PMN (C’+PMN). Background killing <5% is achieved with heat-inactivated C’ (HI C’) or PMN + HI C’. (**B**) Pre-immune, normal (NFS) or immune foal sera obtained on the day of challenge with *R. equi* representing animals with low, medium (Med) or high titers to PNAG mediate killing of intracellular *R. equi* along with C’ and PMN. (**C**) Measurement of percent cytotoxicity by LDH release shows MAb to PNAG or PNAG-immune sera plus C’ and PMN mediate lysis of infected cells. (**D**) Opsonic killing of intracellular *R. equi* requires recognition of cell surface PNAG. Treatment of infected macrophage cultures with dispersin B to digest surface PNAG eliminates killing whereas treatment with the control enzyme, chitinase, has no effect on opsonic killing. Bars represent means of 4-6 technical replicates. Depicted data are representative of 2-3 independent experiments. Bars showing <0% kill represent data wherein the cfu counts were greater than the control of PNAG MAb + PMN + HI C’.

PNAG can be digested with the enzyme dispersin B that specifically recognizes the β-1→6-linked *N*-acetyl glucosamine residues (*38, 39*) but is unaffected by chitinase, which degrades the β-1→4-linked *N*-acetyl glucosamines in chitin. Thus, we treated human macrophages infected for 24 h with *R. equi* with either dispersin B or chitinase to determine if the presence of surface PNAG was critical for killing of intracellular bacteria. Dispersin B treatment markedly reduced the presence of PNAG on the infected cell surface (fig. S7) as well as killing of intracellular *R. equi* by antibody, complement, and PMN (Fig. 4D). Chitinase treatment had no effect on PNAG expression (fig. S7) or killing, indicating a critical role for PNAG intercalated into the macrophage membrane for antibody-mediated killing of intracellular *R. equi*.

### Antibody to PNAG mediates intracellular killing of other intracellular pathogens

To show that antibody to PNAG, complement, and PMN represent a general mechanism for killing of disparate intracellular pathogenic bacteria that express PNAG, we used the above-described system of infected human macrophages to test killing of *M. avium, S. aureus, Neisseria gonorrhoeae*, *L. monocytogenes* and *Bordetella pertussis* by the human MAb to PNAG or horse serum from a foal protected from *R. equi* pneumonia. Human MDM infected with these organisms expressed PNAG on the surface that was not detectable after treatment with dispersin B (fig. S7). When present intracellularly, all of these organisms were killed in the presence of MAb to PNAG or anti-PNAG immune horse serum, complement, and PMN following treatment of the infected cells with the control enzyme, chitinase, but killing was markedly reduced in infected cells treated with dispersin B (Figs. 5A and S8). As with *R. equi*, maximal lysis of infected cells occurred when antibody to PNAG plus complement and PMN were present (Fig. 5B and fig. S9), although when analyzing data from all 5 of these experiments combined there was a modest but significant release of LDH release with antibody to PNAG and complement alone (Fig. 5B and fig. S9).

**Fig. 5.**
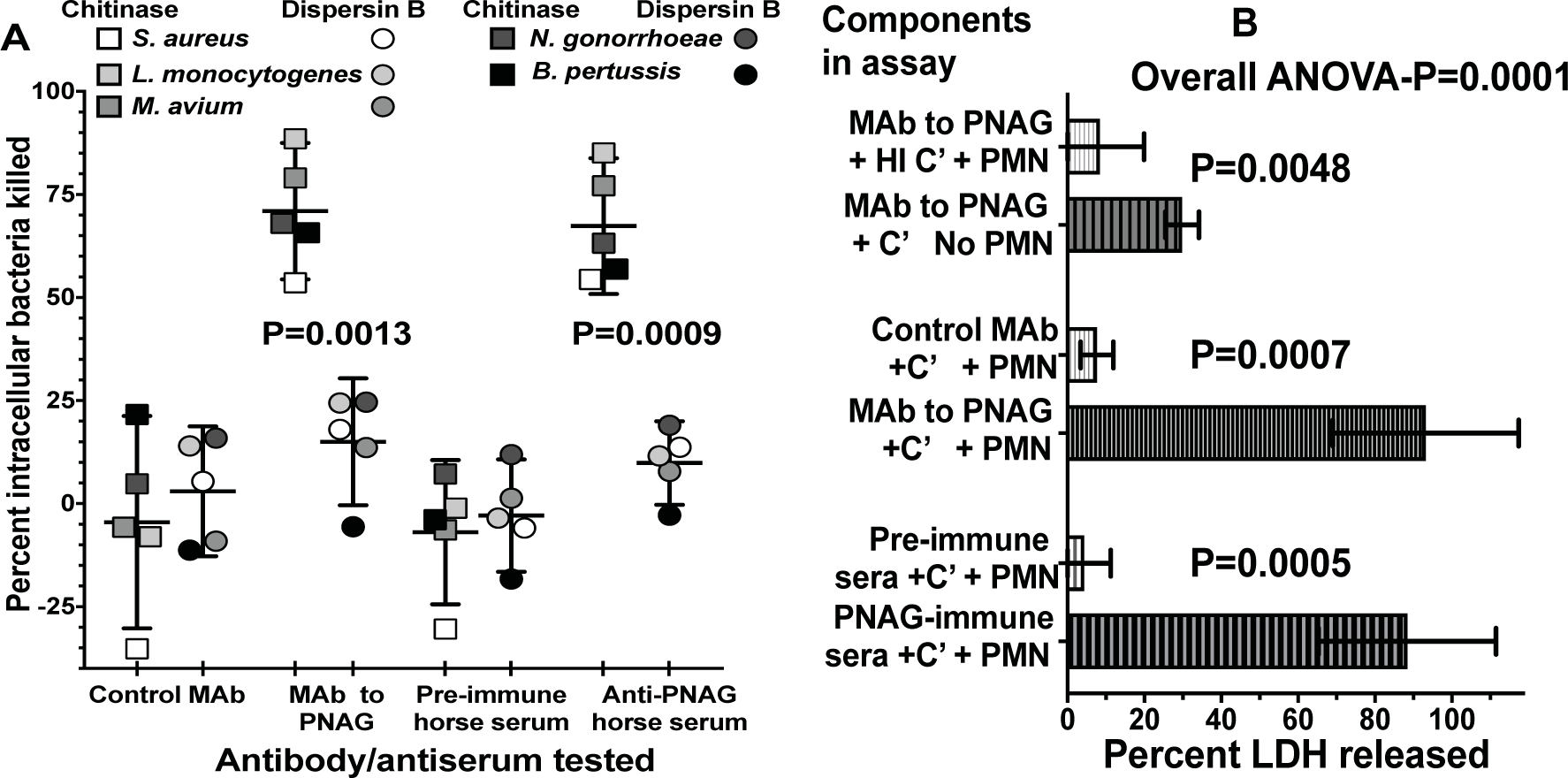
Opsonic killing of multiple intracellular pathogens by antibody to PNAG, complement (C’) and PMN depends on infected-cell surface expression of PNAG and is associated with release of LDH. (A) Killing of 5 different intracellular bacterial pathogens by monoclonal or polyclonal antibody (10% concentration) to PNAG plus PMN and C’ was markedly reduced following treatment of infected cells with Dispersin B (circles) to digest surface PNAG compared with treatment with the control enzyme, Chitinase (squares). Symbols represent indicated bacterial target strain. Horizontal bars represent means, error bars the 95% C.I. Symbols showing <0% kill represent data wherein the cfu counts were greater than the control of PNAG MAb + PMN + HI C’. P values: paired t-tests comparing percent intracellular bacteria killed with each antibody/antiserum tested after Chitinase or Dispersin B treatment. (**B**) Opsonic killing is associated with maximal LDH release from infected cells in the presence of antibody to PNAG, C’ and PMN. Bars represent means from 5 different intracellular pathogens, error bars the 95% C.I., overall ANOVA P value by one-way repeated measures ANOVA, pair wise comparisons determined by two-stage linear step-up procedure of Benjamini, Krieger and Yekutieli.

### Maternal PNAG vaccination and antibody transfer to foals enhances *in vitro* cell-mediated immune responses against *R. equi*

Cell-mediated immune (CMI) responses in vaccinated and unvaccinated, control foals were assessed by detecting production of IFN-γ from peripheral blood mononuclear cells (PBMC) stimulated with a lysate of virulent *R. equi*. IFN-γ production at 2 days of age was significantly (P < 0.05; linear mixed-effects modeling) lower than levels at all other days for both the control and vaccinated groups (Fig. 6A, P value not on graph). There was no difference in IFN-γ production between vaccinated and control foals at day 2 of age. Vaccinated foals had significantly higher (~10-fold) production of IFN-γ in response to *R. equi* stimulation (Fig. 6A) from cells obtained just prior to challenge on days 25-28 of life compared to unvaccinated controls. Subsequent to infection, the controls likely made a CMI response to *R equi*, removing differences days between vaccinates and controls in PBMC IFN-γ production by the age of 56.

**Fig. 6.**
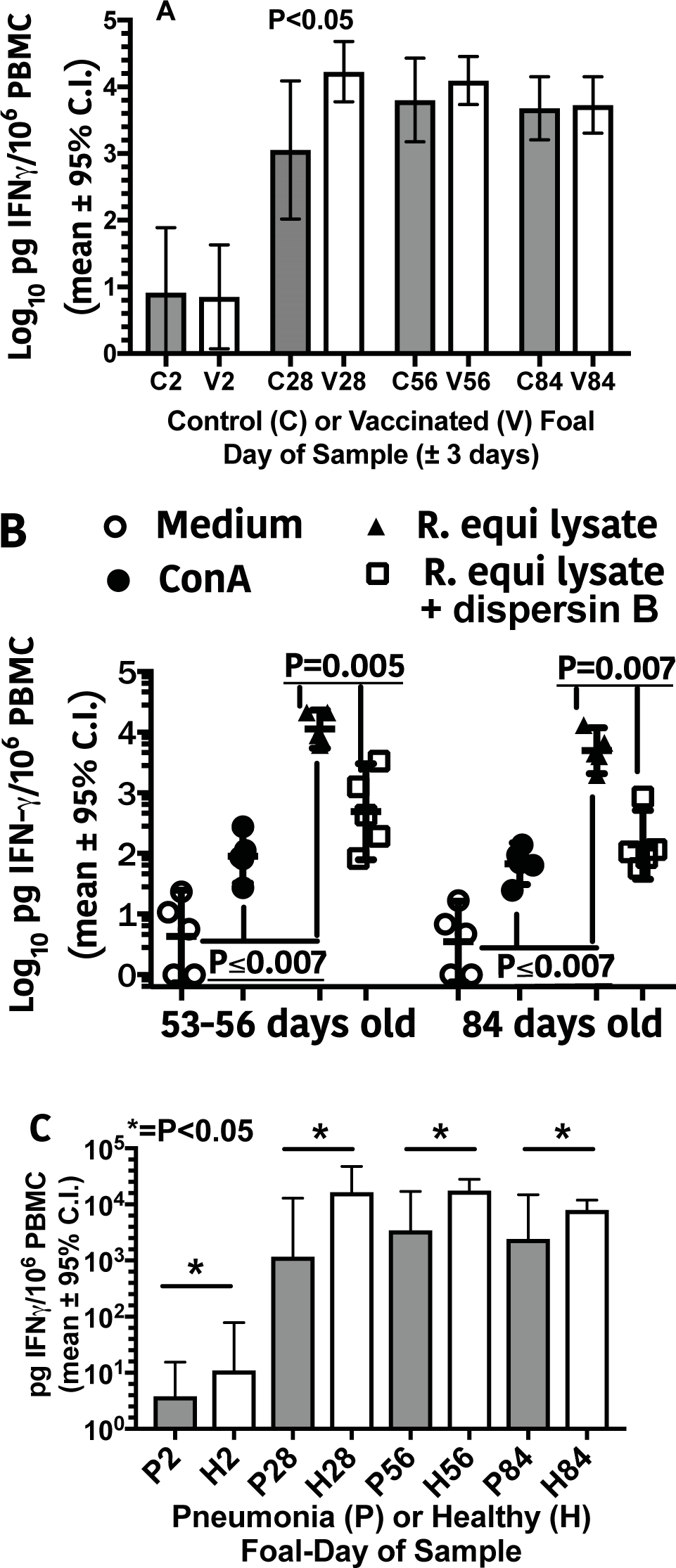
Cell-mediated immune responses of foal PBMCs. **(A)** Foals (N=7 controls, 12 vaccinated) from vaccinated mares (V) had significantly (P <0.05; linear-mixed effects modeling) higher concentrations of IFN-γ produced at 28 days of age (prior to challenge) than control (C) foals in response to stimulation by a lysate of *R. equi*. IFN-γ production at 2 days of age was significantly (P <0.05; linear mixed-effects modeling) lower than those at all other days for both the control and vaccine groups (P values not shown on graph). (**B**) IFN-γ production from PBMC from 5 foals at 56 and 84 days of age following intrabronchial infection with virulent *R. equi.* Stimuli included media only (negative control), Concanavilin A (ConA; positive control), lysate of virulent *R. equi* strain used to infect the foals (*R. equi* Lysate); and the same lysate treated with dispersin B to digest PNAG. All 3 stimulated groups were significantly different from the control at both day 56 and 84 (Overall ANOVA for repeated measures (P < 0.0001); P ≤ 0.0070 for all pairwise comparisons to media only and for pairwise comparison for *R. equi* lysate vs. lysate plus dispersin B (indicated on top of graph), Holm-Sidak’s multiple comparisons test. (**C**) Foals (N=7 controls, 12 vaccinated) that developed pneumonia (P) had significantly (P < 0.05; linear-mixed effects modeling) lower concentrations of IFN-γ expression at each day relative to foals that remained healthy (H).

To substantiate the specificity of this CMI reaction, we demonstrated that stimulation of horse PBMC from vaccinated foals with an *R. equi* lysate treated with the enzyme dispersin B diminished IFN-γ responses by ~90% (Fig. 6B). We also made a *post hoc* comparison of CMI responses between foals that remained healthy and foals that developed pneumonia. In this analysis (Fig. 6C), foals that remained healthy (11 vaccinates and 1 control) had significantly (P < 0.05; linear mixed-effects modeling) higher CMI responses at all ages, including age 2 days, than foals that became ill (1 vaccinate and 6 controls), suggesting that both innate and acquired cellular immunity contribute to resistance to *R. equi* pneumonia. Overall, it appears the maternally derived antibody to PNAG sensitizes foal PBMC to recognize the PNAG antigen and release IFN-γ, which is a known effector of immunity to intracellular pathogens.

## Discussion

In this study we showed maternal immunization against the deacetylated glycoform of the conserved microbial surface polysaccharide PNAG induced antibodies that protected 11 of 12 (91%) ~4-week-old foals from challenge with live, virulent *R. equi*. The attack rate in the controls was 86% (6/7). A confirmatory passive transfer study similarly showed antibody to PNAG infused into foal blood on the day of birth protected 5/5 (100%) of recipients against *R. equi* pneumonia following challenge at 4 weeks of life whereas 100% of 4 controls were diagnosed with *R. equi* pneumonia after challenge. Between the colostral and the infusion passive transfer experiments, a total of 16 of 17 foals with vaccine-induced antibody to PNAG were protected against all evidence of clinical pneumonia (Table S1) due to *R. equi* whereas 10 of 11 non-immune controls had a clinical diagnosis of pneumonia (P < 0.0001, Fisher’s exact test; protected fraction 90%, (95% C.I. 59%-98%, Koopman asymptotic score (*34*)). Vaccine-induced antibody to PNAG deposited complement component C1q onto the purified PNAG antigen, mediated opsonic killing of both extracellular and intracellular *R. equi*, and sensitized PBMC from vaccinated foals to release IFN-γ in response to PNAG. The mechanism of killing of intracellular PNAG-expressing microbes was dependent upon surface expression of this antigen, presumably intercalated into the plasma membrane of the infected host cell following release and intracellular transport of microbial surface vesicles (*37*). This mechanism of killing was shown to be applicable to the PNAG-expressing intracellular pathogens *M. avium, L. monocytogenes, S. aureus, B. pertussis*, and *N. gonorrhoeae*. Overall, these results imply a high potential for efficacy of vaccination against PNAG in protecting against human intracellular pathogens, and with the parallels in pathogenesis of *R. equi* in foals and Mtb in humans, further imply a potential for protective efficacy against human TB.

While immunization-challenge studies such as those performed here are often correlative with protective efficacy against infection and disease, such studies can have limitations in their ability to predict efficacy in the natural setting. Bolus challenges provide an acute insult and immunologic stimulus that mobilizes immune effectors and clears infectious organisms, whereas in a field setting, such as natural acquisition of *R. equi* by foals, infection likely occurs early in life with onset of disease signs taking several weeks to months to develop (*6, 7*). Pulmonary acquisition of Mtb followed by a latent period prior to the emergence of disease occurs in humans exposed to this pathogen (*40*). Thus, it cannot be predicted with certainty that the protective efficacy of antibody to PNAG manifest in the setting of acute, bolus challenge will also be effective when a lower infectious inoculum and more insidious course of disease develops. In this setting, activation of immune effectors may not be of sufficient intensity to take advantage of the opsonic killing activity of antibody, thus allowing progression to disease to occur. In the context of acute challenge, we noted that many of the protected vaccinated foals developed small lung lesions after challenge that rapidly resolved and no disease signs were seen. Finding such lesions by routine ultrasound examination of foals that occurs on farms (*41*) might instigate treatment of subclinical pneumonia if equine veterinarians are either unwilling to monitor foals until clinical signs appear or unconvinced that disease would not ensue in vaccinated foals. This approach would obviate the benefit of vaccination.

The use of animal models to predict vaccine efficacy in humans is fraught with uncertainty, even when non-human primates are used (*42-44*). Thus, whether the protective efficacy shown here for *R. equi* in foals is predictive of efficacy against human infection with Mtb is unknown, even though these 2 organisms share significant pathogenic properties. One difference between these microbes is their growth rates, and this may impact the effectiveness of PNAG immunization of humans. *R. equi* grows rapidly in laboratory culture but Mtb slowly. Also, while *R. equi,* Mtb, and other pathogenic mycobacteria are reasonably closely related genetically, there are likely some differences in surface composition and antigens that might impact the ability of antibody to PNAG to kill mycobacteria, thus limiting the utility of this vaccine for human Mtb. However, we did find that the antibody to PNAG could kill a variety of human pathogens inside of infected macrophages, including a rapidly-growing mycobacterial pathogen, dependent on antibody recognition of PNAG antigen intercalated into the infected cell’s plasma membrane. This finding not only provides an explanation for how antibody to PNAG can mediate protective immunity to intracellular pathogens, but also supports the potential of antibody to PNAG for providing broad-based protective immunity to many intracellular pathogens.

The protection studies described here for *R. equi* disease in foals has led to the implementation of a human trial evaluating the impact of infusion of the fully human IgG_1_ MAb to PNAG (*21*) on latent and new onset TB. The MAb has been successfully tested for safety, pharmacokinetic, and pharmacodynamic properties in a human phase I test (*45*). The trial in TB patients began in September 2017 (South African Clinical Trials Register). The MAb was chosen for initial evaluation to avoid issues of variable immunogenicity that might arise if a vaccine were tried in a TB-infected population, and to have a greater margin of safety in case of untoward effects of immunity to PNAG in the human setting. It is expected the half-life of the MAb will lead to its reduction to pre-infusion levels over 9 to 15 months whereas this might not be the case following vaccination. A successful effect of the MAb on treatment or disease course in TB will lead to an evaluation of immunogenicity and efficacy of a PNAG targeting vaccine in this patient population. The vaccine used here in horse mares was part of a batch of material produced for human phase 1 safety and immunogenicity testing (ClinicalTrials.gov Identifier: NCT02853617), wherein early results indicate that among a small number of vaccinates there were no serious adverse events and high titers of functional antibody elicited in 7 of 8 volunteers given either 75 µg or 150 µg doses twice 28 days apart.

Numerous investigators have studied how antibodies can mediate protection against intracellular bacterial pathogens like TB (*16, 19, 46*), although specific mechanisms of immunity are not well defined. The *in vitro* results we derived indicated that a cell infected with a PNAG-producing pathogen has prominent surface display of this antigen that serves as a target for antibody, complement and PMN to lyse the infected cell and release the intracellular organisms for subsequent opsonic killing. Likely other bacterial antigens are displayed on the infected host cell as well, and thus this system could be used to evaluate the protective efficacy and mechanism of killing of antibodies to other antigens produced by intracellular organisms. Although we have not investigated the basis for the appearance of PNAG in the plasma membrane of infected host cells, we suspect that microbial extracellular vesicles, known to be released by many microbes including Mtb (*47*), are a likely source of the plasma membrane antigen due to trafficking from infected cellular compartments (*37*).

A notable component of the immune response in the foals was the association of release of IFN-γ from PBMC in response to a *R. equi* cell lysate with the protective efficacy of the maternally derived antibody. The response to the lysate significantly dropped after treatment of the lysate with the PNAG-degrading enzyme dispersin B, indicating that an antibody-dependent cellular response to PNAG underlay the IFN-γ response. As this cytokine is well known to be an important component of resistance to human TB (*13*), it was notable that the maternal immunization strategy led to an IFN-γ response. It also appears that the reliance on traditional T-cell effectors recognizing MHC-restricted microbial antigens to provide components of cellular immunity can potentially be bypassed by an antibody-dependent mechanism of cellular responses, further emphasizing how antibody can provide immunity to intracellular pathogens.

This study addressed many important issues related to vaccine development, including the utility of maternal immunization to provide protection against an intracellular pathogen via colostrum to immunologically immature offspring, the efficacy and mechanism of action of antibody to PNAG in protective efficacy, and identification of a role for antibody-dependent IFN-γ release in the response to immunization that likely contributed to full immunity to challenge. The success of immunization in protecting against *R. equi* challenge in foals targeting the broadly synthesized PNAG antigen raises the possibility that this single vaccine could engender protection against many microbial pathogens. While the potential to protect against multiple microbial targets is encouraging, the findings do raise issues as to whether antibody to PNAG will be protective against many microbes or potentially manifest some toxicities or unanticipated enhancements of infection caused by some organisms. Thus, continued monitoring and collection of safety data among animals and humans vaccinated against PNAG is paramount until the safety profile of antibody to PNAG becomes firmly established. Overall, the protective efficacy study in foals against *R. equi* has initiated the pathway to development of PNAG as a vaccine for significant human and animal pathogens, and barring unacceptable toxicity, the ability to raise protective antibodies to PNAG with the 5GlcNH_2_-TT conjugate vaccine portends effective vaccination against a very broad range of microbial pathogens.

## Materials and Methods

### Experimental Design

The objective of the research was to test the ability of maternal vaccination of horse mares with a conjugate vaccine targeting the PNAG antigen to deliver, via colostral transfer, antibody to their offspring that would prevent disease due to intrabronchial *R. equi* challenge at ~4 weeks of life. A confirmatory study using passive infusion of immune or control horse plasma to foals in the first 24 hours of life was also undertaken. The main research subjects were the foals; the secondary subjects were the mares and their immune responses. The experimental design was a randomized, controlled, experimental immunization-challenge trial in horses, with pregnant mares and their foals randomly assigned to the vaccine or control group. Group assignment was made using a randomized, block design for each year. Data were obtained and processed randomly then pooled after unblinding for analysis. Investigators with the responsibility for clinical diagnosis were blinded to the immune status of the foals. An unblinded investigator monitored the data collected to ascertain lack of efficacy and stopping of the infections if 5 or more vaccinated foals developed pneumonia. A similar design was used for the transfusion/passive infusion study, except for the stopping rule.

### Samples size determination

The sample size for the foal protection study was based on prior experience with this model ^6, 30, 48^ indicating a dose of 10^6^ CFU of *R. equi* delivered in half-portions to the left and right lungs via intrabronchial instillation would cause disease in ~85% of foals. Thus, a control group of 7 foals, anticipating 6 illnesses, and a vaccinated group of 12 foals, would have the ability to detect a significant effect at a P value of <0.05 if 75% of vaccinated foals were disease-free using a 1-sided Fisher’s exact test. A 2-sided test is not feasible as one cannot realistically measure a disease rate in vaccinated foals significantly greater than 85%, and any failure to reduce the disease rate would not be indicative of vaccine efficacy. Thus, lack of reduction in disease would lead to rejection of the hypothesis that vaccination against PNAG is effective in preventing *R. equi* pneumonia. Similar criteria were applied to the passive infusion/protection study. All clinical and immunological data to be collected were defined prior to the trial in mares and foals, and no outliers were excluded from the analysis. The primary endpoint was development of clinical *R. equi* pneumonia as defined under Clinical Monitoring below. Experiments were performed over 3 foaling seasons: 2015 and 2016 for the active immunization of pregnant mares, with results from the 2 years of study combined, and 2017 for the passive infusion study.

### Ethics statement

All procedures for this study were reviewed and approved by the Texas A&M Institutional Animal Care and Use Committee (protocol number AUP# IACUC 2014-0374 and IACUC 2016-0233) and the University Institutional Biosafety Committee (permit number IBC2014-112). The foals used in this study were university-owned, and permission for their use was provided in compliance with the Institutional Animal Care and Use Committee procedures. No foals died or were euthanized as a result of this study.

### Vaccine

Mares in the vaccine group received 125 µg (during 2015) or 200 µg (2016) of synthetic pentamers of β 1→6-linked glucosamine conjugated to tetanus toxoid (ratio of oligosaccharide to protein 35-39:1; AV0328, Alopexx Enterprises, LLC, Concord, MA) diluted to 900 µl in sterile medical grade physiological (*i.e.*, 0.9% NaCl) saline solution (PSS) combined with 100 µl of Specol (Stimune^®^ Immunogenic Adjuvant, Prionics, Lelystad, Netherlands, now part of Thermo-Fischer Scientific), a water-in-oil adjuvant. The rationale for increasing the dose in 2016 was that some vaccinated mares had relatively low titers, although all foals of mares in 2015 were protected. Mares in the unvaccinated group were sham injected with an equivalent volume (1 ml) of sterile PSS. All pregnant mares were vaccinated/sham vaccinated 6 and 3 weeks prior to their estimated due dates. For the transfusion of hyperimmune plasma, adult horses (not pregnant) were immunized as above, blood obtained, and hyperimmune plasma produced from the blood by the standard commercial techniques used by Mg Biologics, Ames, Iowa for horse plasma products. Controls received commercially available normal equine plasma prepared from a pool of healthy horses.

### Study populations and experimental infection

Twenty healthy Quarter Horse mare/foal pairs were initially included in this study; 1 unvaccinated mare and her foal were excluded when the foal was stillborn. The unvaccinated group consisted of 7 mare/foal pairs (n = 4 in 2015 and n = 3 in 2016) and the vaccinated group consisted of 12 mare/foal pairs (n = 5 in 2015 and n = 7 in 2016). For the passive infusion of hyperimmune plasma, 9 foals were used, 4 infused with 2 L of commercial normal horse plasma (Immunoglo Serial 1700, Mg Biologics, Ames, IA, USA) and 5 were infused with 2 L of PNAG-hyperimmune plasma produced using standard methods by Mg Biologics. Group assignment was made using a randomized, block design for each year. All foals were healthy at birth and had total serum IgG concentrations >800 mg/dl at 48 h of life using the SNAP^®^ Foal IgG test (IDEXX, Inc., Westbrook, Maine, USA), and remained healthy through the day of experimental challenge. Immediately prior to experimental infection with *R. equi*, each foal’s lungs were evaluated by thoracic auscultation and thoracic ultrasonography to document absence of evidence of pre-existing lung disease.

To study vaccine efficacy, foals were experimentally infected with 1 x 10^6^ of live *R. equi* strain EIDL 5–331 (a virulent, *vapA*-gene-positive isolate recovered from a pneumonic foal). This strain was streaked onto a brain-heart infusion (BHI) agar plate (Bacto Brain Heart Infusion, BD, Becton, Dickinson and Company, Sparks, MD, USA). One CFU was incubated overnight at 37°C in 50 ml of BHI broth on an orbital shaker at approximately 240 rpm. The bacterial cells were washed 3 times with 1 X phosphate-buffered saline (PBS) by centrifugation for 10 min, 3000 x g at 4°C. The final washed pellet was resuspended in 40 ml of sterile medical grade PBS to a final concentration of 2.5 x 10^4^ CFU/ml, yielding a total CFU count of 1 x 10^6^ in 40 ml. Half of this challenge dose (20 ml with 5 x 10^5^) was administered transendoscopically to the left mainstem bronchus and the other half (20 ml with 5 x 10^5^) was administered to the right mainstem bronchus. Approximately 200 µl of challenge dose was saved to confirm the concentration (dose) administered, and to verify virulence of the isolate using multiplex PCR (23).

For transendoscopic infection, foals were sedated using intravenous (IV) injection of romifidine (0.8 mg/kg; Sedivet, Boehringer-Ingelheim Vetmedica, Inc., St. Joseph, MO, USA) and IV butorphanol (0.02 mg/kg; Zoetis, Florham Park, New Jersey, USA). An aseptically-prepared video-endoscope with outer diameter of 9-mm was inserted via the nares into the trachea and passed to the bifurcation of the main-stem bronchus. A 40-mL suspension of virulent EIDL 5–331 *R. equi* containing approximately 1 x 10^6^ viable bacteria was administered transendoscopically, with 20 ml infused into the right mainstem bronchus and 20 ml into the left mainstem bronchus via a sterilized silastic tube inserted into the endoscope channel. The silastic tube was flushed twice with 20 ml of air after each 20-ml bacterial infusion. Foals and their mares were housed individually and separately from other mare and foal pairs following experimental infection.

### Sample collections from mares and foals

Colostrum was collected (approx. 15 ml) within 8 hours of foaling. Blood samples were collected from immunized mares 6 weeks and 3 weeks before their predicted dates of foaling, and on the day of foaling. Blood samples from 4 non-vaccinated mares in the 2015 study were only collected on the day of foaling, whereas blood was collected from the 3 non-vaccinated mares in the 2016 study at the same time-points as those for vaccinated mares. Blood for preparation of hyperimmune plasma was collected from immunized adult horses 2 weeks after the second injection of 200 µg of the 5GlcNH_2_-TT vaccine in 0.1 ml of Specol.

Blood samples were drawn from foals on day 2 (the day after foaling), and at 4, 6, 8, and 12 weeks of age. Samples at 4 weeks (25-28 days of life) were collected prior to infection. Blood was collected in EDTA tubes for complete blood count (CBC) testing, in lithium heparinized tubes for PBMC isolation, and in clot tubes for serum collection.

Transendoscopic tracheobronchial aspiration (T-TBA) was performed at the time of onset of clinical signs for any foals developing pneumonia and at age 12 weeks for all foals (end of study) by washing the tracheobronchial tree with sterile PBS solution delivered through a triple-lumen, double-guarded sterile tubing system (MILA International, Inc. Erlanger, KY, USA).

### Clinical monitoring

From birth until the day prior to infection, foals were observed twice daily for signs of disease. Beginning the day prior to infection, rectal temperature, heart rate, respiratory rate, signs of increased respiratory effort (abdominal lift, flaring nostrils), presence of abnormal lung sounds (crackles or wheezes, evaluated for both hemithoraces), coughing, signs of depressed attitude (subjective evidence of increased recumbence, lethargy, reluctance to rise), and nasal discharges were monitored and results recorded twice daily through 12 weeks (end of study). Thoracic ultrasonography was performed weekly to identify evidence of peripheral pulmonary consolidation or abscess formation consistent with *R. equi* pneumonia. Foals were considered to have pneumonia if they demonstrated ≥3 of the following clinical signs: coughing at rest; depressed attitude (reluctance to rise, lethargic attitude, increased recumbency); rectal temperature >39.4°C; respiratory rate ≥60 breaths/min; or, increased respiratory effort (manifested by abdominal lift and nostril flaring). Foals were diagnosed with *R. equi* pneumonia if they had ultrasonographic evidence of pulmonary abscessation or consolidation with a maximal diameter of ≥2.0 cm, positive culture of *R. equi* from T-TBA fluid, and cytologic evidence of septic pneumonia from T-TBA fluid. The primary outcome was the proportion of foals diagnosed with *R. equi* pneumonia. Secondary outcomes included the duration of days meeting the case definition, and the sum of the total maximum diameter (TMD) of ultrasonography lesions over the study period. The TMD was determined by summing the maximum diameters of each lesion recorded in the 4^th^ to the 17^th^ intercostal spaces from each foal at every examination; the sum of the TMDs incorporates both the duration and severity of lesions. Foals diagnosed with *R. equi* pneumonia were treated with a combination of clarithromycin (7.5 mg/kg; PO; q 12 hour) and rifampin (7.5 mg/kg; PO; q 12 hour) until both clinical signs and thoracic ultrasonography lesions had resolved. Foals also were treated as deemed necessary by attending veterinarians (AIB; NDC) with flunixin meglumine (0.6 to 1.1 mg/kg; PO; q 12-24 hour) for inflammation and fever.

### Immunoglobulin ELISAs

Systemic humoral responses were assessed among foals by indirectly quantifying concentrations in serum by ELISA from absorbance values of PNAG-specific total IgG and by IgG subisotypes IgG_1_, IgG_4/7_, and IgG_3/5_. ELISA plates (Maxisorp, Nalge Nunc International, Rochester, NY, USA) were coated with 0.6 µg/ml of purified PNAG ^49^ diluted in sensitization buffer (0.04M PO_4_, pH 7.2) overnight at 4°C. Plates were washed 3 times with PBS with 0.05% Tween 20, blocked with 120 µl PBS containing 1% skim milk for 1 hour at 37°C, and washed again. Mare and foal serum samples were added at 100 µl in duplicate to the ELISA plate and incubated for 1 hour at 37°C. Serum samples were initially diluted in incubation buffer (PBS with 1% skim milk and 0.05% Tween 20) to 1:100 for total IgG titers, 1:64 for IgG_1_ and IgG_4/7_ detection, and to 1:256 for IgG_3/5_ detection. A positive control from a horse previously immunized with the 5GlcNH_2_-TT vaccine and known to have a high titer, along with normal horse serum known to have a low titer, were included in each assay for total IgG titers. For the subisotype assays, immune rabbit serum (rabbit anti-5GLcNH_2_-TT) was diluted to a concentration of 1:102,400 as a positive control and used as the denominator to calculate the endpoint OD ratio of the experimental OD values. The immune rabbit serum was used to account for inter-plate variability and negative control of normal rabbit serum were included with the equine serum samples. After 1 hour incubation at 37°C, the plates were washed 3 times as described above. For total IgG titers, rabbit anti-horse IgG whole molecule conjugated to alkaline phosphatase (Sigma-Aldrich, St. Louis, MO, USA) was used to detect binding. For IgG subisotype detection, 100 µl of goat-anti-horse IgG_4/7_ (Lifespan Biosciences, Seattle, WA, diluted at 1:90,000), or goat anti-horse IgG_3/5_ (Bethyl Laboratories, Montgomery, TX, USA, diluted at 1:30,000) conjugated to horseradish peroxidase, or mouse anti-horse IgG_1_ (AbD Serotec, Raleigh, NC, USA), diluted at 1:25,000) were added to the wells and incubated for 1 hour at room temperature. For the IgG_1_ subisotype, goat antibody to mouse IgG (Bio-Rad, Oxford, England, diluted at 1:1000) conjugated to peroxidase was used for detection. Plates were washed again, and for the total IgG titers pNPP substrate (1 mg/ml) was added while for peroxidase-conjugated antibody to mouse IgG, SureBlue Reserve One Component TMB Microwell Peroxidase Substrate (SeraCare, Gaithersburg, MD, USA) was added to the wells. Plates were incubated for 15 to 60 minutes at 22°C in the dark. The reaction was stopped by adding sulfuric acid solution to the wells. Optical densities were determined at 450 nm by using microplate readers. Equine subisotype concentrations of PNAG-specific IgG_1_, IgG_4/7_, and IgG_3/5_ were also quantified in colostrum of each mare using the same protocol described above for serum. Colostral samples were diluted in incubation buffer (PBS with 1% skim milk and 0.05% Tween 20) to 1:8,192 for IgG_1_, 1:4096 for IgG_4/7_ detection, and at 1:64 for IgG_3/5_ detection. For total IgG endpoint titers were calculated by linear regression using a final OD_405_nm value of 0.5 to determine the reciprocal of the maximal serum dilution reaching this value. For IgG subisotypes, an endpoint OD titer was calculated by dividing the experimental OD values with that achieved by the positive control on the same plate.

### PNAG expression by clinical isolates of *R. equi*

Clinical isolates of *R. equi* were obtained from the culture collection at the Equine Infectious Disease Laboratory, Texas A&M University College of Veterinary Medicine & Biomedical Sciences. All strains were originally isolated from foals diagnosed with R. *equi* pneumonia and were obtained from geographically distinct locations. *R. equi* strains were grown overnight on BHI agar then swabbed directly onto glass slides, air dried and fixed by exposure for 1 min to methanol at 4°C. Samples were labeled with either 5 µl of a 5.2 µg/ml concentration of MAb F598 to PNAG directly conjugated to Alexa Fluor 488 or control MAb F429 to alginate, also directly conjugated to Alexa Fluor 488, for 4 hours at room temperature. During the last 5 min of this incubation, 500 nM of Syto83 in 0.5% BSA/PBS pH 7.4 was added to stain nucleic acids (red fluorophore). Samples were washed and mounted for immunofluorescent microscopic examination as described ^11^.

### Analysis of PNAG expression in infected horse tissues and human monocyte-derived macrophage cultures

Paraffinized sections of lungs obtained at necropsy from foals with *R. equi* pneumonia were provided by the Texas A&M College of Veterinary Medicine & Biomedical Sciences histopathology laboratory. Slides were deparaffinized using EzDewax and blocked overnight at 4C with 0.5% BSA in PBS. Samples were washed then incubated with the fluorophore-conjugated MAb F598 to PNAG or control MAb F429 to alginate described above for 4 hours at room temperature. Simultaneously added was a 1:500 dilution (in BSA/PBS) of a mouse antibody to the virulence associated Protein A of *R. equi*. Binding of the mouse antibody to *R. equi* was detected with a donkey antibody to mouse IgG conjugated to Alexa Fluor 555 at a dilution of 1:250 in BSA/PBS. Samples were washed and mounted for immunofluorescence microscopic examination.

To detect PNAG expression in cultured human monocyte-derived macrophages (MDM), prepared as described below in opsonic killing assays, the infected MDM were washed and fixed with 4% paraformaldehyde in PBS for 1 hour at room temperature. To visualize PNAG on the surface of infected cells, MDM cultures were incubated with the fluorophore-conjugated MAb F598 to PNAG or control MAb F429 to alginate for 4-6 hours at room temperature. Samples were then imaged by confocal microscopy to visualize extracellular PNAG expression. Next, these same samples were treated with 100% methanol at 4°C for 5 min at room temperature to permeabilize the plasma membrane. Samples were washed with PBS then incubated with either 5.2 µg/ml of MAb F598 to PNAG or MAb F429 to alginate for 1-2 hours at room temperature, washed in PBS then a 1:300 dilution in PBS of donkey antibody to human IgG labeled with Alexa Fluor 555 added for 4-6 hours at room temperature. Samples were washed and mounted for immunofluorescence microscopic examination.

### C1q deposition assays

Serum endpoint titers for deposition of equine complement component C1q onto purified PNAG were determined by ELISA. ELISA plates were sensitized with PNAG and blocked with skim milk as described above, dilutions of different horse sera added in 50 µl-volumes after which 50 µl of 10% intact, normal horse serum was added. After 60 minutes incubation at 37°C, plates were washed and 100 µl of goat anti-human C1q, which also binds to equine C1q, diluted 1:1,000 in incubation buffer added and plates incubated at room temperature for 60 minutes. After washing, 100 µl of rabbit anti-goat IgG whole molecule conjugated to alkaline phosphatase and diluted 1:1,000 in incubation buffer was added and a 1-hour incubation at room temperature carried out. Washing and developing of the color indicator was then carried out as described above, and endpoint titers determined as described above for IgG titers by ELISA.

### Opsonic killing assays

To determine opsonic killing of *R. equi,* bacterial cultures were routinely grown overnight at 37°C on chocolate-agar plates, then killing assessed using modifications of previously described protocols ^49^. Modifications included use of EasySep™ Human *Neutrophil Isolation* Kits (Stem Cell Technologies Inc., Cambridge, Massachusetts, USA) to purify PMN from blood, and use of gelatin-veronal buffer supplemented with Mg^++^ and Ca^++^ (Boston Bioproducts, Ashland, Massachusetts, USA) as the diluent for all assay components. Final assay tubes contained, in a 400-µl volume, 2 × 10^5^ human PMN, 10% (final concentration) *R. equi*-absorbed horse serum as a complement source, 2 × 10^5^ *R equi* cells and the serum dilutions. Tubes were incubated with end-over-end rotation for 90 minutes then diluted in BHI with 0.05% Tween and plated for bacterial enumeration.

For intracellular opsonic killing assays, human monocytes were isolated from peripheral blood using the EasySep™ Direct Human Monocyte Isolation Kit (Stem Cell Technologies) and 2 × 10^4^ cells placed in a 150 µl volume of RPMI and 10% heat-inactivated autologous human serum in flat-bottom 96-well tissue culture plates for 5-6 days with incubation at 37°C in 5% CO_2_. Differentiated cells were washed and 5 × 10^5^ CFU of *R. equi* in RPMI and 10% heat-inactivated autologous human serum added for 30 minutes. Next, cells were washed and 150 µl of RPMI plus10% autologous serum with 50 µg gentamicin sulfate/ml added and cells incubated for 24 hours at 37°C in 5% CO_2_. For some experiments, 50 µl of 400 µg/ml of either chitinase (Sigma-Aldrich) or dispersin B (Kane Biotech, Winnipeg, Manitoba), a PNAG-degrading enzyme ^37, 50^, dissolved in Tris-buffered saline, pH 6.5, were added directly to gentamicin containing wells and plates incubated for 2 hours at 37°C in 5% CO_2_. Cell cultures were washed then combinations of 50 µl of MAb or foal serum, 50 µl of 30% *R. equi*-absorbed human serum as a complement source, or heat-inactivated complement as a control, and 50 µl containing 1.5 × 10^5^ human PMN, isolated as described above, added. Controls lacked PMN or had heat-inactivated complement used in place of active complement, and final volumes made up with 50 µl of RPMI 1640 medium. After a 90-minute incubation at 37°C in 5% CO_2_, 10 µl samples were taken from selected wells for analysis of lysis by lactate dehydrogenase release, and 100 µl of trypsin/EDTA with 0.1% Triton X100 added to all wells lyse the cells via a 10-minute incubation at 37°C. Supernatants were diluted and plated on chocolate agar for bacterial enumeration as described above.

### Cell-mediated immunity

The cell-mediated immune response to vaccination was assessed by measuring IFN-γ production from isolated horse PBMCs that were stimulated with an *R. equi* antigen lysate of strain EIDL 5–331, or the same lysate digested for 24 hours at 37°C with 100 µg/ml of dispersin B. The PBMCs were isolated using a Ficoll-Paque gradient separation (GE Healthcare, Piscataway, NJ, USA) and resuspended in 1X RPMI-1640 media with L-glutamine (Gibco, Life Technologies, Grand Island, NY, USA), 15% fetal bovine serum (Gibco), and 1.5% penicillin-streptomycin (Gibco). The PBMCs were cultured for 48 hours at 37°C in 5% CO_2_ with either media only, the mitogen Concanavalin A (positive control; 2.5 µg/ml, Sigma-Aldrich), or *R. equi* lysate representing a multiplicity of infection of 10. After 48 hours, supernatants from each group were harvested and frozen at −80°C until examined for IFN-γ production using an equine IFNγ ELISA kit (Mabtech AB, Nacka Strand, Stockholm, Sweden) according to the manufacturer’s instructions. Optical densities were determined using a microplate reader and standard curves generated to determine IFN-γ concentrations in each sample using the Gen 5 software (Biotek, Winooski, VT, USA).

### Statistical methods

Categorical variables with independent observations were compared using chi-squared or, when values for expected cells were ≤5, Fisher’s exact tests. For estimation of the 95% C.I. of the relative risk, the Koopman asymptotic score (*34*) was determined.

Continuous, independent variables were compared between 2 groups using either paired t-tests or Mann-Whitney tests and between > 2 groups using the Kruskal-Wallis test with pairwise *post hoc* comparisons made using Dunn’s procedure. Continuous variables with non-independent observations (*i.e.*, repeated measures) were compared using linear mixed-effects modeling with an exchangeable correlation structure and individual mare or foal as a random effect. Survival times were compared non-parametrically using the log-rank test. All analyses were performed using S-PLUS statistical software (Version 8.2, TIBCO, Inc., Seattle, Wash, USA) or the PRISM 7 statistical program. Mixed-effect model fits were assessed using diagnostic residual plots and data were transformed (log_10_) when necessary to meet distributional assumptions of modeling; post hoc pairwise comparisons among levels of a variable (*e.g.*, age) were made using the method of Sidak (*48*). Significance was set at P ≤ 0.05 and adjustment for multiple comparisons made.

## Funding

This study was funded by a grant from the Morris Animal Foundation (MAF) and Alopexx Vaccine LLC. The results have not been reviewed or endorsed by MAF, and the views expressed do not necessarily reflect the views of the MAF, its officers, directors, affiliates or agents. Additional support was provided by the Link Equine Research Endowment, Texas A&M University. We thank Dr. Deborah Hung from the Broad Institute of MIT and Harvard and the Massachusetts General Hospital for provision of GFP-expressing *M. tuberculosis* cells for analysis.

## Author contributions

C.C-B., J.N.R., A.I.B., M.V., S.R.,T.Z., S.G. G.B.P. and N.D.C. designed and executed experiments. C.C-B., J.N.R., A.I.B., D.V., N.D.C., and G.B.P. analyzed data. J.N.R., A.I.B., and N.D.C. conducted the animal immunization and challenge studies. N.D.C. and G.P. conducted the statistical analysis. C.C-B., J.N.R., A.I.B., D.V., N.D.C. and G.B.P. wrote and edited the manuscript.

## Conflict of interest

Gerald B. Pier is an inventor of intellectual properties [human monoclonal antibody to PNAG and PNAG vaccines] that are licensed by Brigham and Women’s Hospital to Alopexx Vaccine, LLC, and Alopexx Pharmaceuticals, LLC, entities in which GBP also holds equity. As an inventor of intellectual properties, GBP also has the right to receive a share of licensing-related income (royalties, fees) through Brigham and Women’s Hospital from Alopexx Pharmaceuticals, LLC, and Alopexx Vaccine, LLC. GBP’s interests were reviewed and are managed by the Brigham and Women’s Hospital and Partners Healthcare in accordance with their conflict of interest policies.

Colette Cywes-Bentley is an inventor of intellectual properties [use of human monoclonal antibody to PNAG and use of PNAG vaccines] that are licensed by Brigham and Women’s Hospital to Alopexx Pharmaceuticals, LLC. As an inventor of intellectual properties, CC-B also has the right to receive a share of licensing-related income (royalties, fees) through Brigham and Women’s Hospital from Alopexx Pharmaceuticals, LLC.

Noah D. Cohen has received an unrestricted gift to the EIDL from Alopexx Vaccines, LLC.

Daniel Vlock, holds an equity share and potential royalty income from Alopexx Vaccines, LLC for vaccines to PNAG and monoclonal antibody to PNAG from Alopexx Pharmaceuticals, LLC.

## Data and materials availability

Materials used in this study including vaccines, serum samples, and microbial strains may be obtained through an MTA with either Brigham and Women’s Hospital or Texas A&M College of Veterinary Medicine & Biomedical Sciences.

---

**Table S1.**
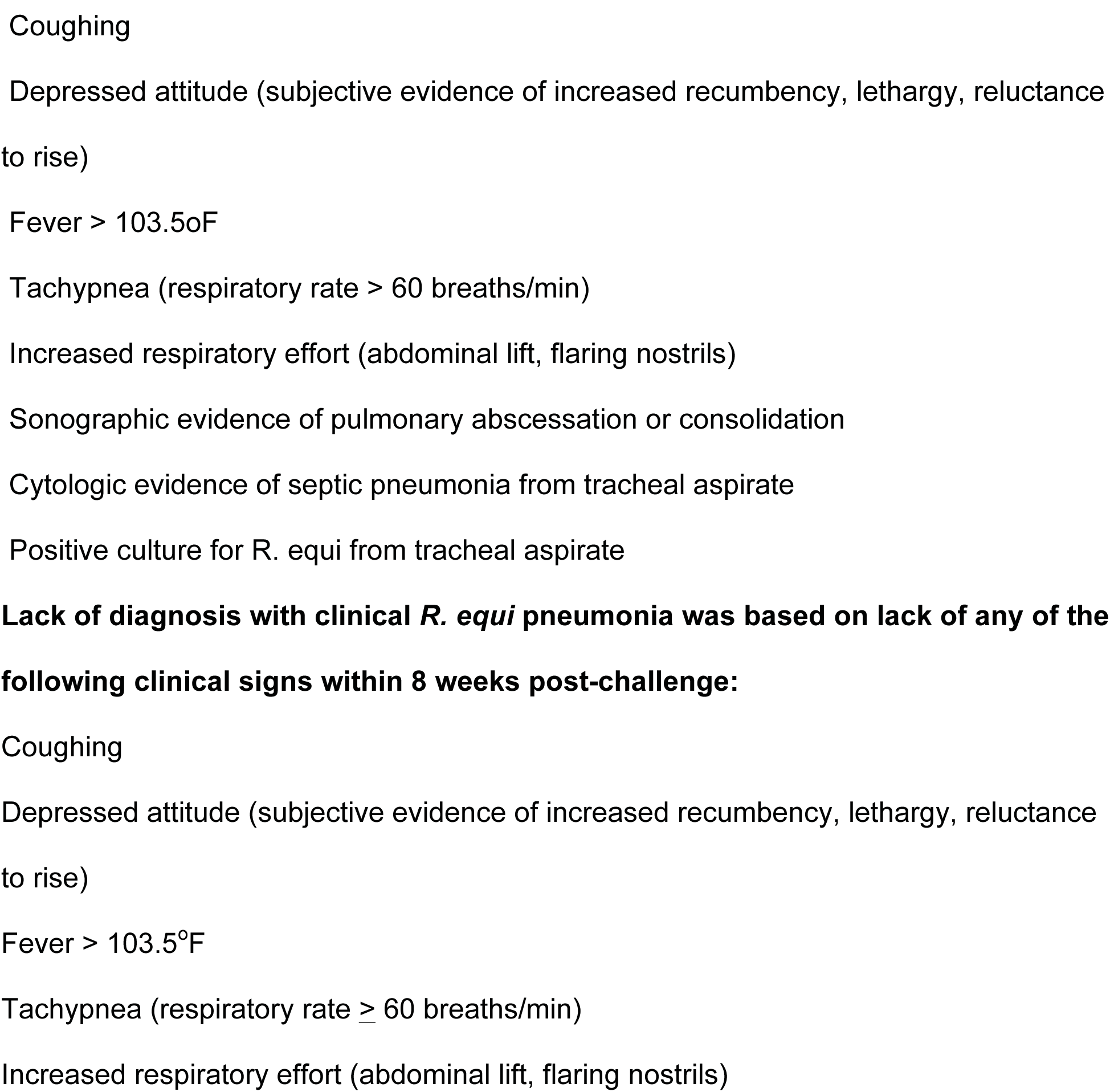
Case definition for diagnosis of *R. equi* clinical pneumonia Cases had all of the following clinical signs within 3 weeks of challenge.

**Table S2.** Duration of clinical signs in foals from vaccinated or unvaccinated mares.

| Variable | Mare Unvaccinated<br>(n=7) | Mare Vaccinated<br>(n=12) | P value<sup>a</sup> |
| --- | --- | --- | --- |
| Days meeting case<br>definition (range) | 10<sup>b</sup> (0 to 26) | 0 (0 to 41) | 0.0046 |
| Days from first to last day<br>meeting case definition (range) | 20 (0 to 32) | 0 (0 to 48) | 0.0046 |
| Days temperature > 103.0°F<br>(range) | 8 (1 to 32) | 2 (0 to 12) | 0.0263 |
| Days temperature > 103.0°F<br>and coughing (range) | 4 (0 to 8) | 0 (0 to 13) | 0.0220 |

<sup>a</sup>P values derived using the Wilcoxon rank-sum test.
<sup>b</sup>Median

**Table S3.** Duration of clinical signs in foals infused with control or PNAG-hyperimmune plasma.

| Variable | Standard plasma<br>(n=4) | PNAG plasma<br>(n=5) | P<br>value<sup>a</sup> |
| --- | --- | --- | --- |
| Days meeting case<br>definition (range) | 10.5<sup>b</sup> (1 to 14) | 0 (0 for all) | 0.0108 |
| Days from first to last day<br>meeting case definition (range) | 11.5 (1 to 14) | 0 (0 for all) | 0.0104 |
| Days temperature > 103.0°F<br>(range) | 11.5 (1 to 15) | 2 (0 to 4) | 0.0811 |
| Days temperature > 103.0°F<br>and coughing (range) | 9.5 (0 to 11) | 0 (0 to 3) | 0.0072 |
| Duration of ultrasound lesions<br>in weeks (range) | 2 (1 to 4) | 0 (0 to 3) | 0.2029 |
| Maximum of TMD<sup>c</sup> (cm) | 10.1 (4.0 to 29.8) | 0 (0 to 3.4) | 0.0179 |
| Sum of TMDs (cm) | 13.2 (4.0 to 50.5) | 0 (0 to 7.9) | 0.0342 |

<sup>a</sup>P values derived using the Wilcoxon rank-sum test.
<sup>b</sup>Median
<sup>c</sup>TMD = total (sum) of maximum diameters of lesions observed on a given day

**Fig. S1.**
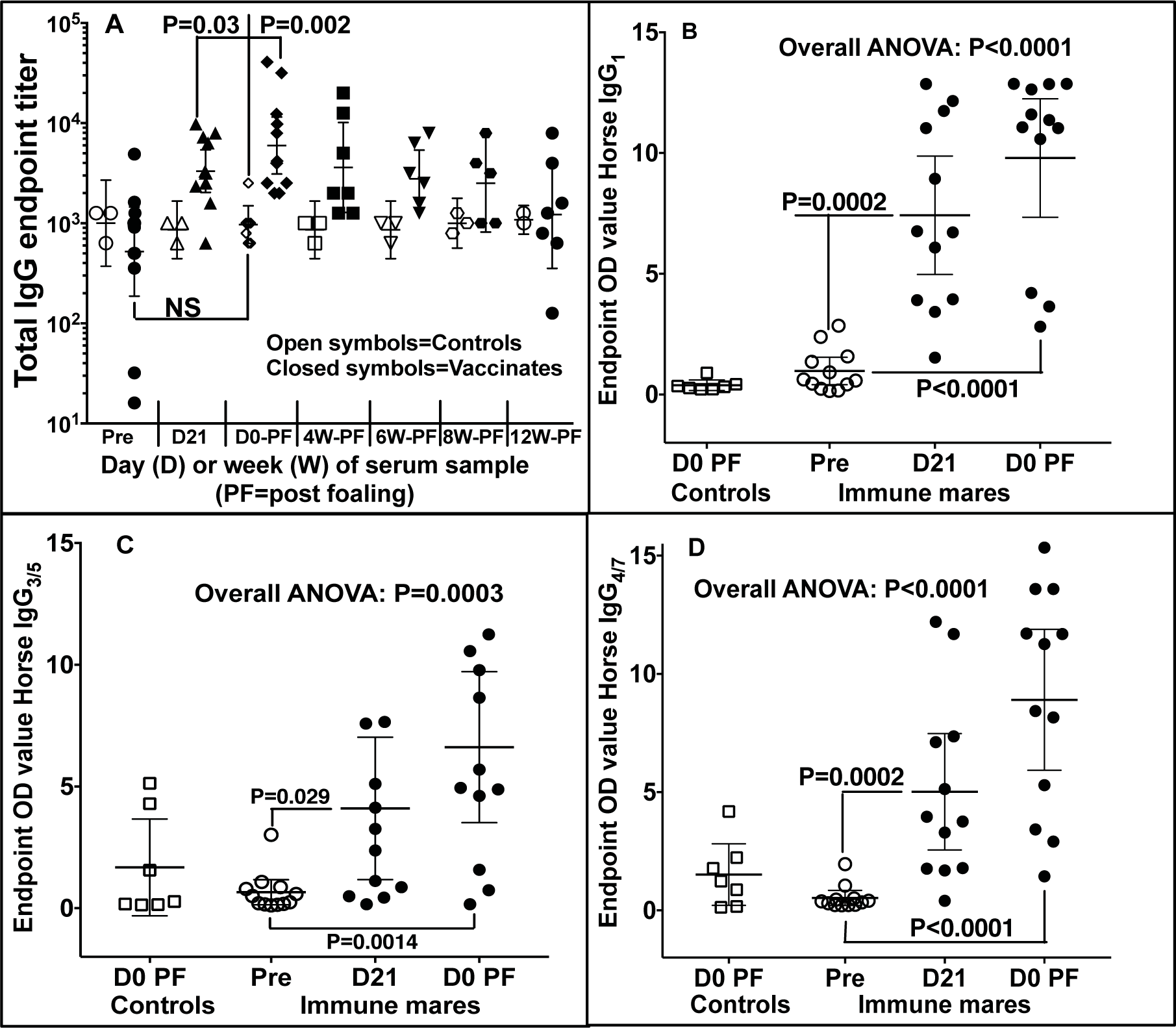
Serum IgG and IgG subisotype titers to PNAG in immunized mare sera. Serum end-point titers of IgG or IgG subisotypes are plotted by vaccine group as a function of age in days. (**A**) Total IgG antibody end-point titers to PNAG were significantly higher in sera of immunized mares at D21 and D0 PF compared with titers in sera of control mares at D0 PF. There was no significant (NS) difference in the IgG titers of the vaccinated mares at pre-immunization and controls at D0 PF. (**B-D**) Concentrations of IgG_1_, IgG_4/7_, and IgG_3/5_ were significantly higher in mares in the vaccinated group at D21 and D0 PF as indicated on the figure. Statistical comparisons were made by linear mixed-effects regression with exchangeable correlation structure, using the mare as random effect (to account for repeated measures) and multiple comparisons determined by the method of Sidak.

**Fig. S2.**
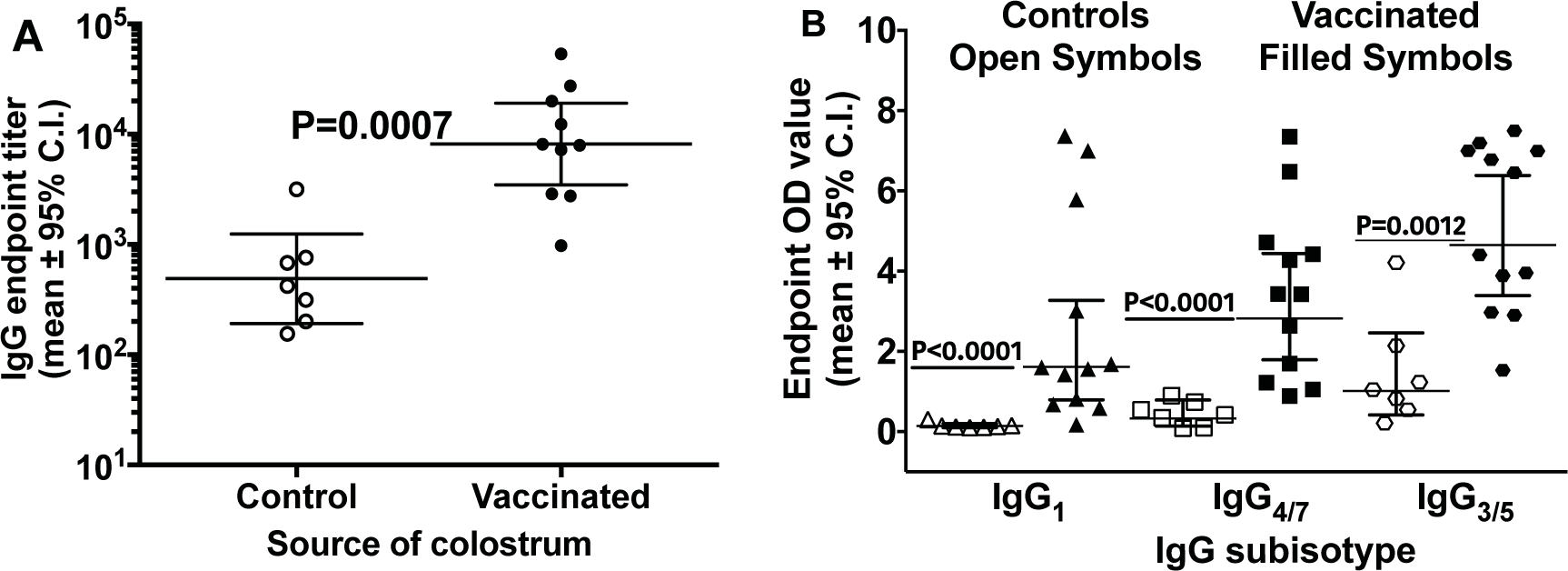
IgG and IgG subisotype titers in mare colostra on day of foaling. End-point colostral titers of IgG or IgG subisotype. End-point values are plotted by vaccine group. (**A**) Total IgG antibody end-point titers to PNAG were significantly higher in colostra of vaccinated mares (N=10, 2 samples not tested due to limited quantities) compared with colostra of control mares N=7). (**B**) Concentrations of IgG_1_, IgG_4/7_, and IgG_3/5_ were significantly higher in colostra of mares in the vaccinated group (N=12). Statistical comparisons were made by the Wilcoxon rank-sum test.

**Fig S3.**
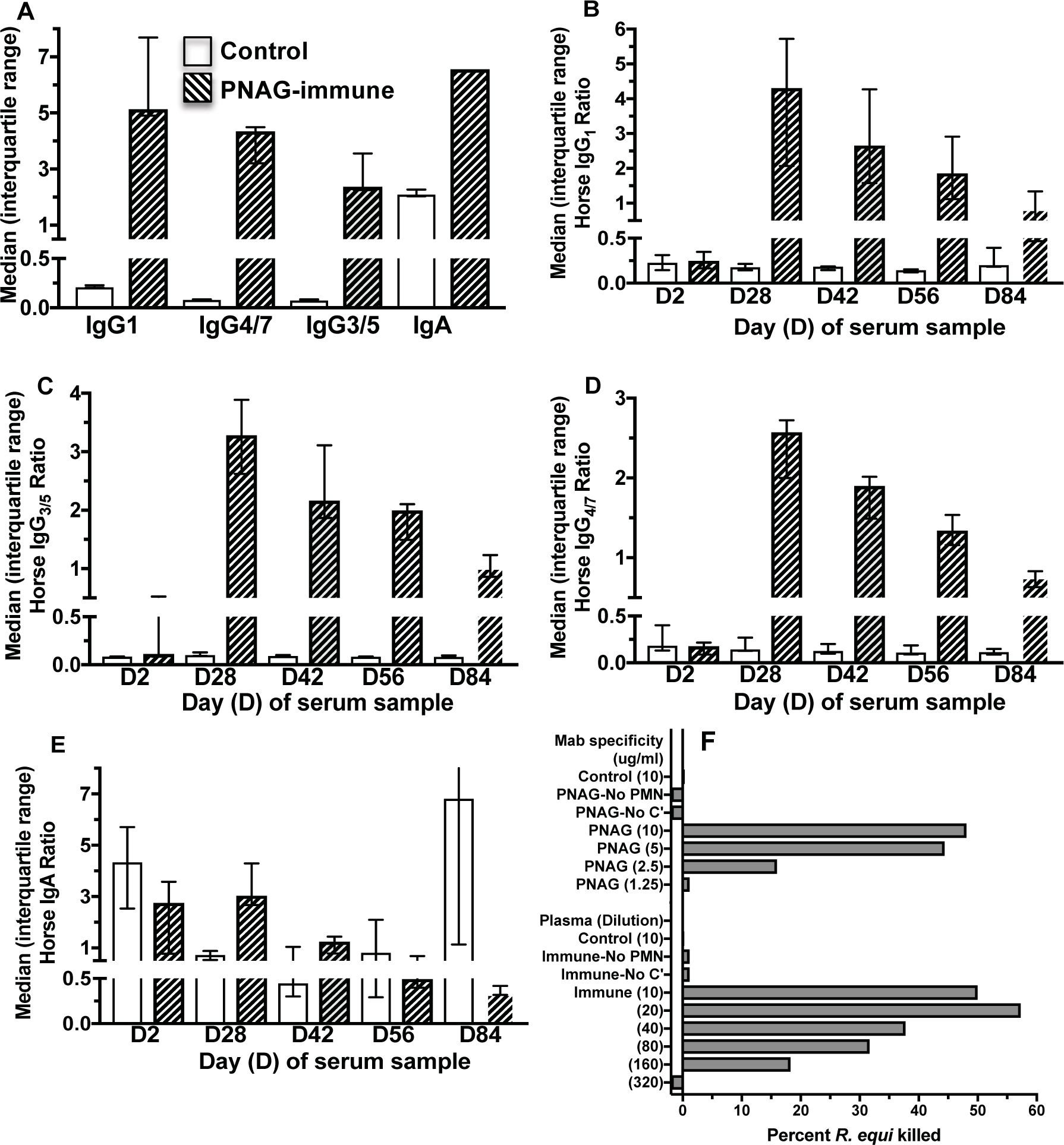
Antibody titers to PNAG in control and hyperimmune horse plasma infused into newborn foals on day 1 of life. (**A**) Titers of IgG subisotypes and IgA in control (open bars) and PNAG-immune (hatched bars) plasmas. (**B**) horse IgG_1_, (**C**) horse, IgG_3/5_, (**D**) horse IgG_4/7_, or (**E**) horse IgA in sera at day indicated on X-axis. Bars represent medians, error bars the interquartile ranges. OD ratios of IgG_1_, IgG_4/7_, and IgG_3/5_ did not differ significantly over time in controls but were significantly (P < 0.05) greater than age 2 days in foals transfused with anti-PNAG plasma at ages 28, 42, and 56 days (IgG_1_), or ages 28, 42, 56, and 84 for IgG_4/7_ and IgG_3/5_. OD ratio of IgA did not differ significantly (P > 0.05) among the different days for anti-PNAG-transfused foals, but controls had significantly (P < 0.05) higher IgA titers at age 2 days compared to control titers on days 28, 42, and 56. The OD ratio values for control foals’ IgA on day 84 was significantly (P < 0.05) greater than those of control foals on days 28, 42, and 56. IgA titers between controls and anti-PNAG-transfused foals differed significantly (P < 0.05) at day 84 only. All P values were determined by mixed-effects linear regression. (**F**) Opsonic killing of *R. equi* EIDL 990 by antibody in control or immune plasma. Monoclonal antibodies (MAb) were used as controls, as were tubes lacking PMN or complement (C’) as indicated. Bars represent means of technical replicates.

**Fig. S4.**
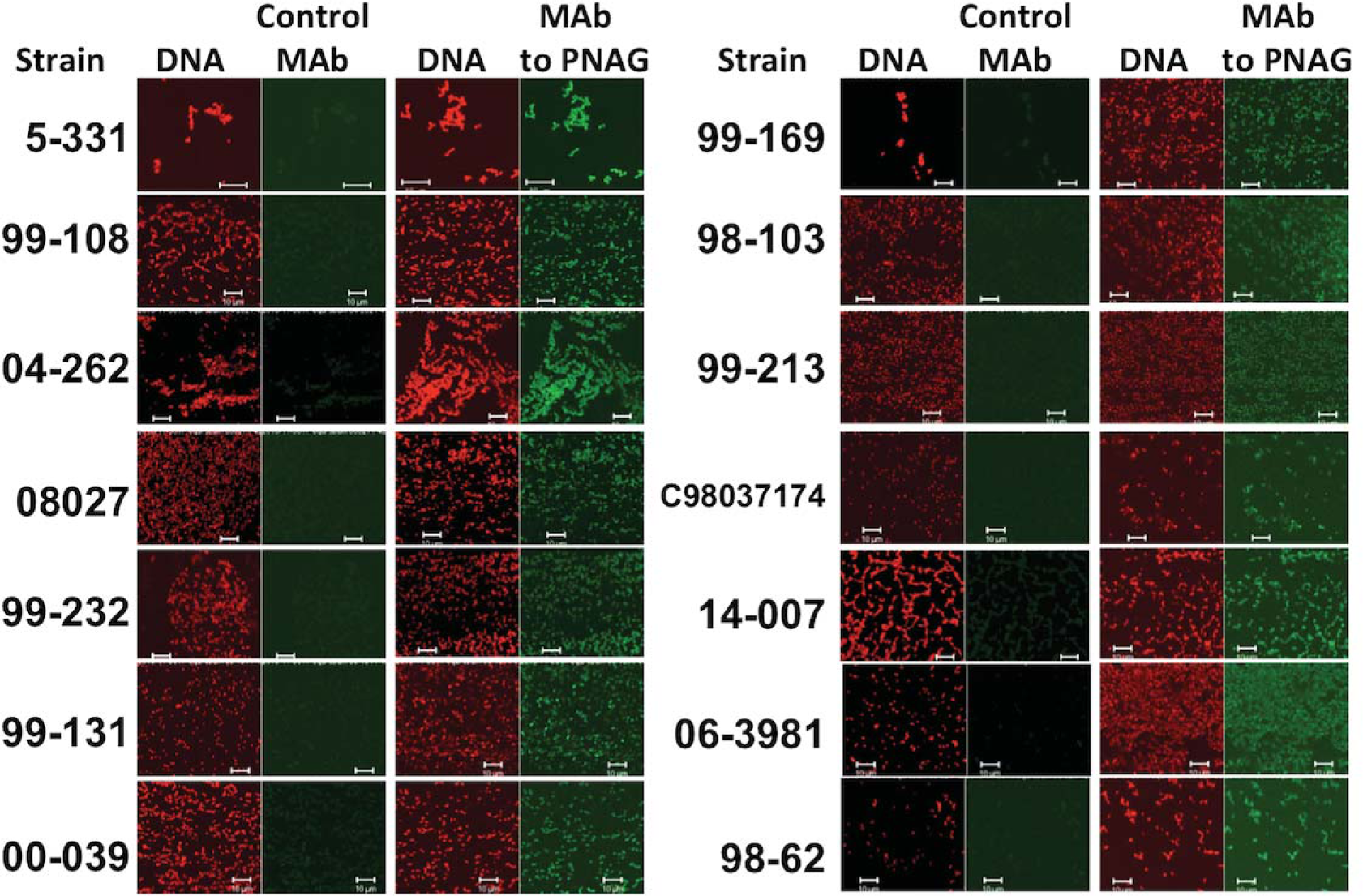
PNAG expression by *R. equi* clinical isolates. Designated individual clinical isolates of R. equi were reacted with either control MAb F429 to *P. aeruginosa* alginate or MAb F598 to PNAG, both directly conjugated to Alexa Fluor 488. Binding to PNAG on bacterial cells was visualized by immunofluorescence microscopy. Left-hand panel in each pair shows DNA visualized by red-fluorophore Syto 83. Right-hand panel in each pair is green if PNAG detected by MAb F598.

**Fig. S5.**
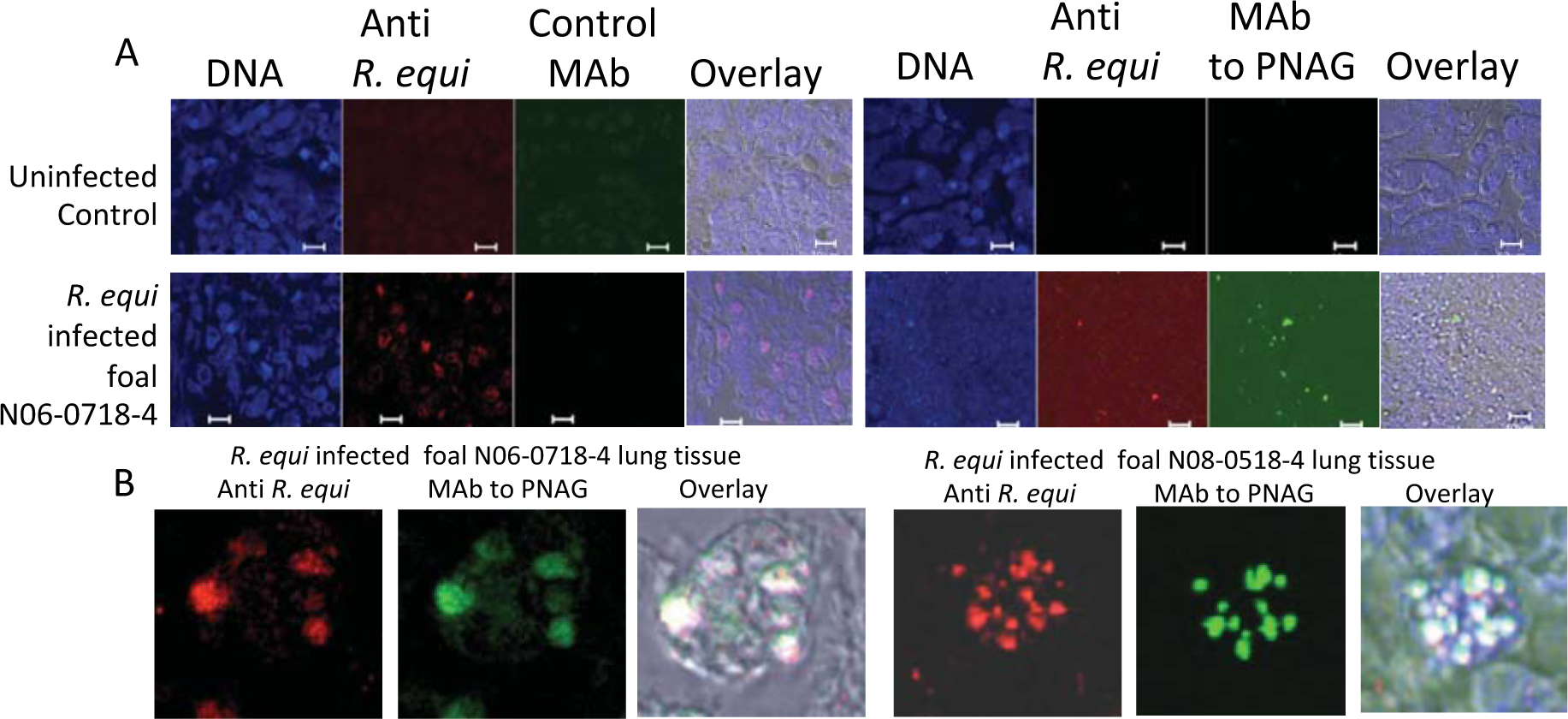
Expression of PNAG in lungs of *R. equi* infected foals. Either an uninfected control lung or lungs from foals with *R. equi* pneumonia were reacted with the indicated antibody to detect the presence of *R. equi* (red, antibody to VapA protein), PNAG (Green, MAb F598) or control MAb F429 to alginate. (**A**) Low power (40X) sections indicating presence of *R. equi* and closely associated PNAG in infected lung. Bars = 10 µm. (**B**) Higher magnification (60 X) shows individual infected cells in 2 different foal lung sections with PNAG-expressing *R. equi* contained in apparent intracellular vesicles.

**Fig. S6.**
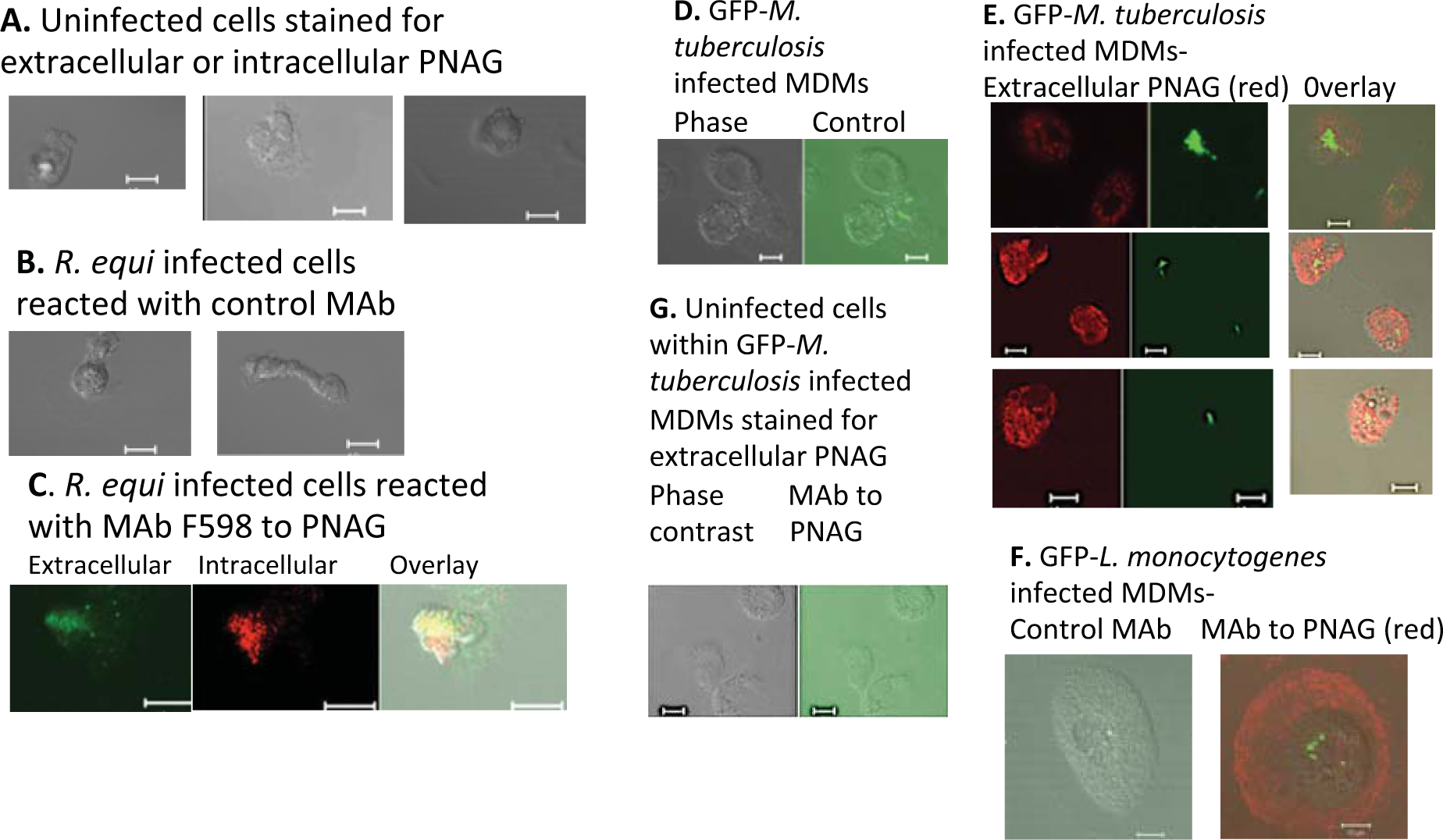
Surface and intracellular expression of PNAG in infected human MDM. Detection of PNAG either on the surface or within the infected cell was determined by first reacting cultures with MAb F598 to PNAG or control MAb F429, both directly conjugated to Alexa Fluor 488 (green fluorophore), on paraformaldehyde-fixed cells then washing and permeabilizing the cells with ice-cold methanol followed by reaction with the MAbs and secondary antibody to human IgG conjugated to Alexa Fluor 555 (red/orange). (**A-G**) Cells and treatments indicated in figure. White bars = 10 µm.

**Fig. S7.**
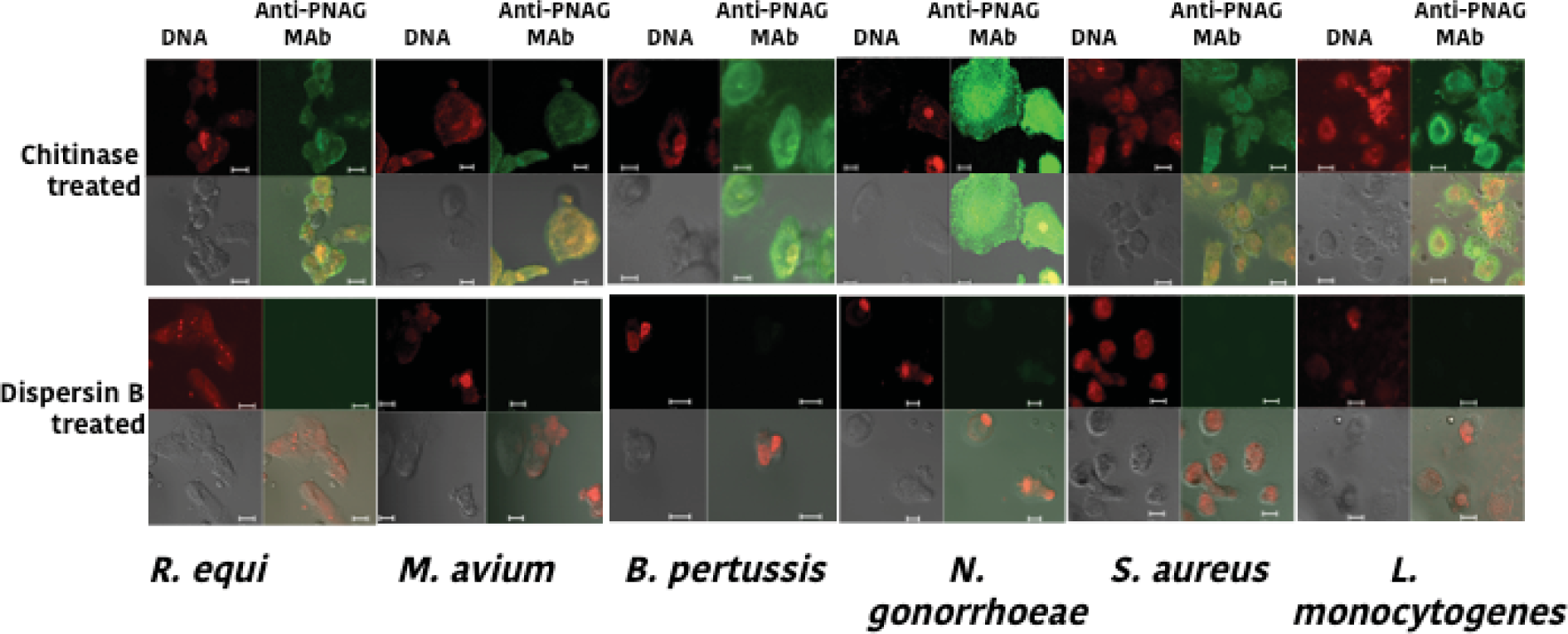
Human MDM cells infected with PNAG-expressing intracellular pathogens have high levels of the PNAG antigen on their surface that is removed by treatment with dispersin B. PNAG on infected cell surfaces was detected by reacting cultures with MAb F598 to PNAG or control MAb F429, both directly conjugated to Alexa Fluor 488 (green fluorophore), on paraformaldehyde-fixed cells. Infected bacterial strains and treatments indicated in figure. For each figure, upper left quadrant shows nuclear DNA (red), upper right quadrant shows PNAG (green if present), lower left quadrant shows phase contrast, lower right quadrant shows co-localization of DNA and PNAG (yellow-green to yellow to orange if present). White bars = 10 µm.

**Fig. S8.**
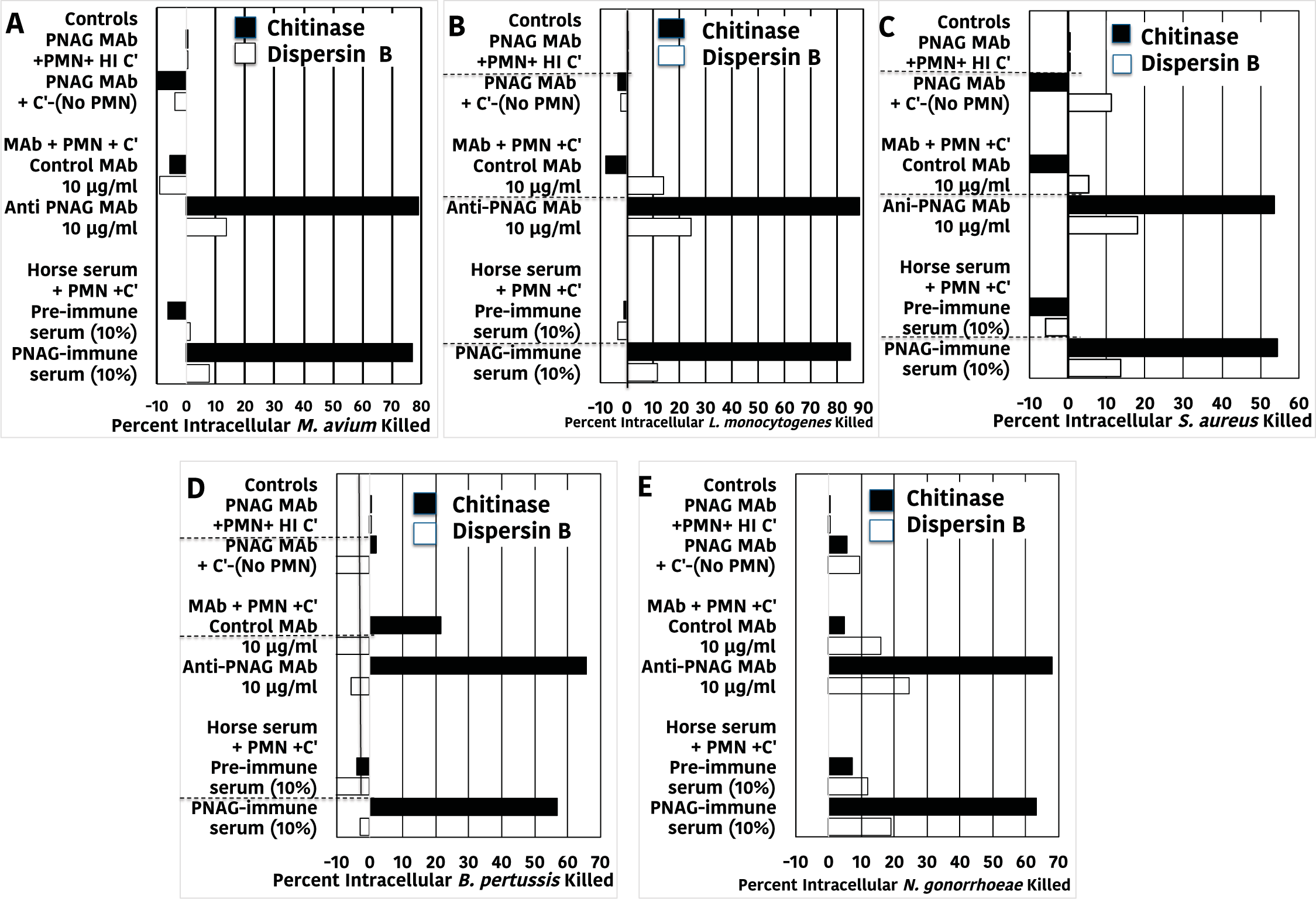
Opsonic killing of intracellular pathogens by antibody to PNAG, complement (C’) and PMN depends on infected-cell surface expression of PNAG. (**A-F**) Killing of intracellular bacteria by antibody, PMN and C’ was markedly reduced following treatment of infected cells with Dispersin B (open bars) to digest surface PNAG compared with treatment with the control enzyme, Chitinase (black bars). Depicted data are representative of 2-3 independent experiments. Bars represent means of 6 technical replicates. Bars showing <0% kill represent data wherein the cfu counts were greater than the control of PNAG MAb + PMN + HI C’.

**Fig. S9.**
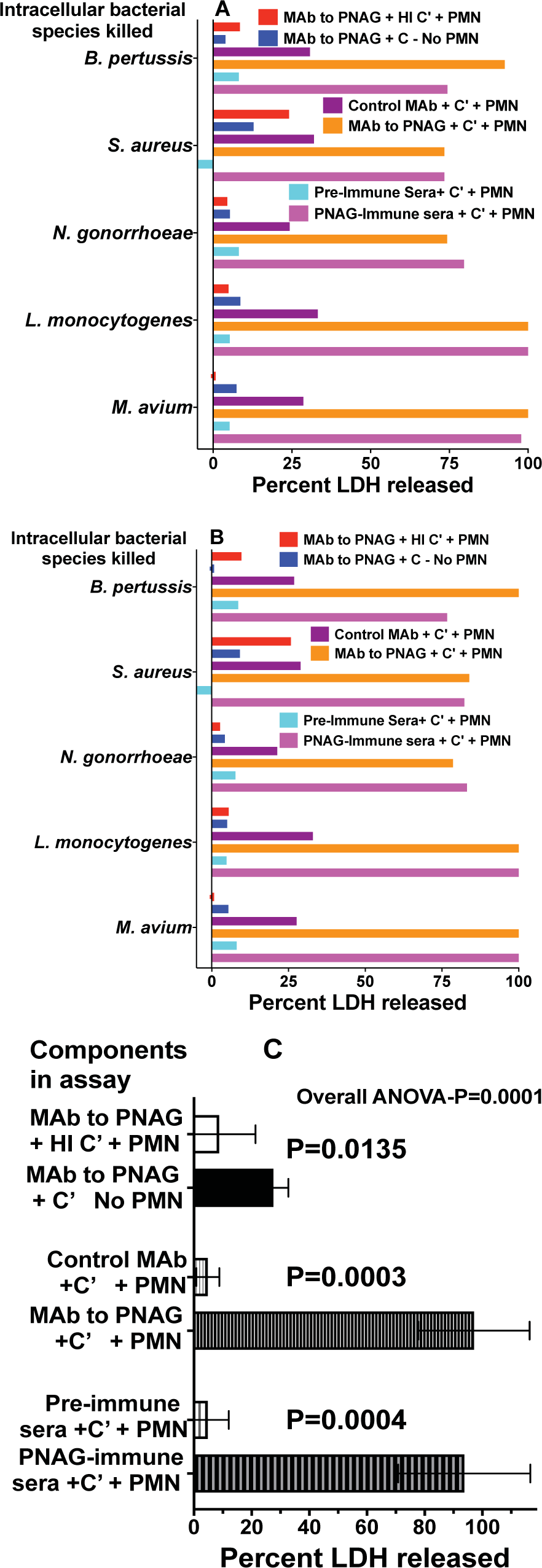
Release of LDH from cells infected with intracellular pathogens following antibody plus immune effector treatment. (**A and B**) Replicate experiments measuring release of LDH from cells infected with indicated pathogen treated with 10 µg/ml control or anti-PNAG monoclonal or 10% polyclonal antibody plus indicated immune effector. Bars indicate means of quadruplicates. (**C**) Summary of LDH release for experiment in figure S9B. Mean (bars) and 95% C.I. (error bars) indicate release of LDH from cells infected with all five intracellular pathogens. Overall ANOVA P value by one-way repeated measures ANOVA, pair wise comparisons determined by two-stage linear step-up procedure of Benjamini, Krieger and Yekutieli.

## References

1. Global Tuberculosis Report 2016. Geneva: World Health Organization (2017). *WHO. Global Tuberculosis Report 2016*. (2017).

2. P. Mendez-Samperio, Global efforts in the development of vaccines for tuberculosis: requirements for improved vaccines against *Mycobacterium tuberculosis*. Scand. J. Immunol. 84, 204–210 (2016).

3. M. K. O’Shea, H. McShane, A review of clinical models for the evaluation of human TB vaccines. Hum. Vaccin. Immunother. 12, 1177–1187 (2016).

4. S. Bhargava, S. Choubey, S. Mishra, Vaccines against tuberculosis: A review. Indian J. Tuberc. 63, 13–18 (2016).

5. T. J. Scriba, S. H. Kaufmann, P. Henri Lambert, M. Sanicas, C. Martin, O. Neyrolles, Vaccination against tuberculosis with whole-cell mycobacterial vaccines. J. Infect. Dis. 214, 659–664 (2016).

6. N. D. Cohen, *Rhodococcus equi* foal pneumonia. Vet. Clin. North Am. Equine Pract. 30, 609–622 (2014).

7. S. Giguere, N. D. Cohen, M. K. Chaffin, S. A. Hines, M. K. Hondalus, J. F. Prescott, N. M. Slovis, Rhodococcus equi: clinical manifestations, virulence, and immunity. J. Vet. Intern. Med. 25, 1221–1230 (2011).

8. S. M. Reuss, N. D. Cohen, Update on bacterial pneumonia in the foal and weanling. Vet. Clin. North Am. Equine Pract. 31, 121–135 (2015).

9. S. Herath, C. Lewis, M. Nisbet, Increasing awareness of *Rhodococcus equi* pulmonary infection in the immunocompetent adult: a rare infection with poor prognosis. N. Z. Med. J. 126, 165–174 (2013).

10. A. V. Yamshchikov, A. Schuetz, G. M. Lyon, *Rhodococcus equi* infection. Lancet Infect. Dis. 10, 350–359 (2010).

11. C. Cywes-Bentley, D. Skurnik, T. Zaidi, D. Roux, R. B. Deoliveira, W. S. Garrett, X. Lu, J. O’Malley, K. Kinzel, T. Zaidi, A. Rey, C. Perrin, R. N. Fichorova, A. K. Kayatani, T. Maira-Litran, M. L. Gening, Y. E. Tsvetkov, N. E. Nifantiev, L. O. Bakaletz, S. I. Pelton, D. T. Golenbock, G. B. Pier, Antibody to a conserved antigenic target is protective against diverse prokaryotic and eukaryotic pathogens. Proc. Natl. Acad. Sci. U. S. A. 110, E2209–2218 (2013).

12. D. Skurnik, C. Cywes-Bentley, G. B. Pier, The exceptionally broad-based potential of active and passive vaccination targeting the conserved microbial surface polysaccharide PNAG. Expert Rev. Vaccines 15, 1041–1053 (2016).

13. I. V. Lyadova, A. V. Panteleev, Th1 and Th17 cells in tuberculosis: protection, pathology, and biomarkers. Mediators Inflamm. 2015, 854507 (2015).

14. M. Raviglione, G. Sulis, Tuberculosis 2015: Burden, challenges and strategy for control and elimination. Infect. Dis. Rep. 8, 6570 (2016).

15. A. D. Harries, D. Maher, S. Graham, World Health Organization., TB/HIV: a clinical manual. (World Health Organization, Geneva, ed. 2nd, 2004), pp. 210 p.

16. R. Prados-Rosales, L. Carreño, T. Cheng, C. Blanc, B. Weinrick, A. Malek, T. L. Lowary, A. Baena, M. Joe, Y. Bai, R. Kalscheuer, A. Batista-Gonzalez, N. A. Saavedra, L. Sampedro, J. Tomás, J. Anguita, S.-C. Hung, A. Tripathi, J. Xu, A. Glatman-Freedman, W. R. Jacobs, Jr., J. Chan, S. A. Porcelli, J. M. Achkar, A. Casadevall, Enhanced control of *Mycobacterium tuberculosis* extrapulmonary dissemination in mice by an arabinomannan-protein conjugate vaccine. PLOS Pathog. 13, e1006250 (2017).

17. H. Li, X. X. Wang, B. Wang, L. Fu, G. Liu, Y. Lu, M. Cao, H. Huang, B. Javid, Latently and uninfected healthcare workers exposed to TB make protective antibodies against *Mycobacterium tuberculosis*. Proc. Natl. Acad. Sci. U S A 114, 5023–5028 (2017).

18. L. L. Lu, A. W. Chung, T. R. Rosebrock, M. Ghebremichael, W. H. Yu, P. S. Grace, M. K. Schoen, F. Tafesse, C. Martin, V. Leung, A. E. Mahan, M. Sips, M. P. Kumar, J. Tedesco, H. Robinson, E. Tkachenko, M. Draghi, K. J. Freedberg, H. Streeck, T. J. Suscovich, D. A. Lauffenburger, B. I. Restrepo, C. Day, S. M. Fortune, G. Alter, A functional role for antibodies in tuberculosis. Cell 167, 433–443 (2016).

19. J. M. Achkar, A. Casadevall, Antibody-mediated immunity against tuberculosis: implications for vaccine development. Cell Host Microbe 13, 250–262 (2013).

20. J. Zuniga, D. Torres-Garcia, T. Santos-Mendoza, T. S. Rodriguez-Reyna, J. Granados, E. J. Yunis, Cellular and humoral mechanisms involved in the control of tuberculosis. Clin. Dev. Immunol. 2012, Article ID: 193923 (2012).

21. C. Kelly-Quintos, L. A. Cavacini, M. R. Posner, D. Goldmann, G. B. Pier, Characterization of the opsonic and protective activity against *Staphylococcus aureus* of fully human monoclonal antibodies specific for the bacterial surface polysaccharide poly-*N*-acetylglucosamine. Infect. Immun. 74, 2742–2750 (2006).

22. C. Kelly-Quintos, A. Kropec, S. Briggs, C. Ordonez, D. A. Goldmann, G. B. Pier, The role of epitope specificity in the human opsonic antibody response to the staphylococcal surface polysaccharide PNAG. J. Infect. Dis. 192, 2012–2019 (2005).

23. M. L. Gening, T. Maira-Litran, A. Kropec, D. Skurnik, M. Grout, Y. E. Tsvetkov, N. E. Nifantiev, G. B. Pier, Synthetic β-(1->6)-linked N-acetylated and nonacetylated oligoglucosamines used to produce conjugate vaccines for bacterial pathogens. Infect. Immun. 78, 764–772 (2010).

24. T. Maira-Litran, A. Kropec, D. Goldmann, G. B. Pier, Biologic properties and vaccine potential of the staphylococcal poly-*N*-acetyl glucosamine surface polysaccharide. Vaccine 22, 872–879 (2004).

25. N. Cerca, K. K. Jefferson, T. Maira-Litran, D. B. Pier, C. Kelly-Quintos, D. A. Goldmann, J. Azeredo, G. B. Pier, Molecular basis for preferential protective efficacy of antibodies directed to the poorly-acetylated form of staphylococcal poly-*N*-acetyl-β-(1-6)-glucosamine. Infect. Immun. 75, 3406–3413 (2007).

26. L. V. Bentancor, J. M. O’Malley, C. Bozkurt-Guzel, G. B. Pier, T. Maira-Litran, Poly-*N*-acetyl-β-(1-6)-glucosamine is a target for protective immunity against *Acinetobacter baumannii* infections. Infect. Immun. 80, 651–656 (2012).

27. D. Skurnik, M. R. Davis, Jr., D. Benedetti, K. L. Moravec, C. Cywes-Bentley, D. Roux, D. C. Traficante, R. L. Walsh, T. Maira-Litran, S. K. Cassidy, C. R. Hermos, T. R. Martin, E. L. Thakkallapalli, S. O. Vargas, A. J. McAdam, T. D. Lieberman, R. Kishony, J. J. Lipuma, G. B. Pier, J. B. Goldberg, G. P. Priebe, Targeting pan-resistant bacteria with antibodies to a broadly conserved surface polysaccharide expressed during infection. J. Infect. Dis. 205, 1709–1718 (2012).

28. M. L. Horowitz, N. D. Cohen, S. Takai, T. Becu, M. K. Chaffin, K. K. Chu, K. G. Magdesian, R. J. Martens, Application of Sartwell’s model (lognormal distribution of incubation periods) to age at onset and age at death of foals with *Rhodococcus equi* pneumonia as evidence of perinatal infection. J. Vet. Intern. Med. 15, 171–175 (2001).

29. M. Sanz, A. Loynachan, L. Sun, A. Oliveira, P. Breheny, D. W. Horohov, The effect of bacterial dose and foal age at challenge on *Rhodococcus equi* infection. Vet. Microbiol. 167, 623–631 (2013).

30. J. N. Rocha, N. D. Cohen, A. I. Bordin, C. N. Brake, S. Giguere, M. C. Coleman, R. C. Alaniz, S. D. Lawhon, W. Mwangi, S. D. Pillai, Oral administration of electron-beam inactivated *Rhodococcus equi* Failed to protect foals against intrabronchial infection with live, virulent R. equi. PLoS One 11, e0148111 (2016).

31. M. J. Flaminio, B. R. Rush, E. G. Davis, K. Hennessy, W. Shuman, M. J. Wilkerson, Characterization of peripheral blood and pulmonary leukocyte function in healthy foals. Vet. Immunol. Immunopathol. 73, 267–285 (2000).

32. T. L. Sturgill, S. Giguere, L. J. Berghaus, D. J. Hurley, M. K. Hondalus, Comparison of antibody and cell-mediated immune responses of foals and adult horses after vaccination with live *Mycobacterium bovis* BCG. Vaccine 32, 1362–1367 (2014).

33. C. Ryan, S. Giguere, Equine neonates have attenuated humoral and cell-mediated immune responses to a killed adjuvanted vaccine compared to adult horses. Clin. Vaccine Immunol. 17, 1896–1902 (2010).

34. P. A. R. Koopman, Confidence intervals for the ratio of two binomial proportions. Biometrics 40, 513–517 (1984).

35. D. Skurnik, M. Merighi, M. Grout, M. Gadjeva, T. Maira-Litran, M. Ericsson, D. A. Goldmann, S. S. Huang, R. Datta, J. C. Lee, G. B. Pier, Animal and human antibodies to distinct *Staphylococcus aureus* antigens mutually neutralize opsonic killing and protection in mice. J. Clin. Invest. 9, 3220–3233 (2010).

36. G. B. Pier, D. Boyer, M. Preston, F. T. Coleman, N. Llosa, S. Mueschenborn-Koglin, C. Theilacker, H. Goldenberg, J. Uchin, G. P. Priebe, M. Grout, M. Posner, L. Cavacini, Human monoclonal antibodies to *Pseudomonas aeruginosa* alginate that protect against infection by both mucoid and nonmucoid strains. J. Immunol. 173, 5671–5678 (2004).

37. W. L. Beatty, H. J. Ullrich, D. G. Russell, Mycobacterial surface moieties are released from infected macrophages by a constitutive exocytic event. Eur. J. Cell Biol. 80, 31–40 (2001).

38. J. E. Kerrigan, C. Ragunath, L. Kandra, G. Gyemant, A. Liptak, L. Janossy, J. B. Kaplan, N. Ramasubbu, Modeling and biochemical analysis of the activity of antibiofilm agent Dispersin B. Acta. Biol. Hung. 59, 439–451 (2008).

39. E. Fazekas, L. Kandra, G. Gyemant, Model for β-1,6-N-acetylglucosamine oligomer hydrolysis catalysed by Dispersin B, a biofilm degrading enzyme. Carb. Res. 363, 7–13 (2012).

40. R. J. Basaraba, R. L. Hunter, Pathology of tuberculosis: How the pathology of human tuberculosis informs and directs animal models. Microbiol. Spectr. 5, (2017).

41. M. Venner, K. Astheimer, M. Lammer, S. Giguere, Efficacy of mass antimicrobial treatment of foals with subclinical pulmonary abscesses associated with *Rhodococcus equi*. J. Vet. Intern. Med. 27, 171–176 (2013).

42. A. J. Hessell, N. L. Haigwood, Animal models in HIV-1 protection and therapy. Curr. Opin. HIV AIDS 10, 170–176 (2015).

43. B. B. Policicchio, I. Pandrea, C. Apetrei, Animal models for HIV cure research. Front. Immunol. 7, 12 (2016).

44. P. J. Cardona, A. Williams, Experimental animal modelling for TB vaccine development. Int. J. Infect. Dis. 56, 268–273 (2017).

45. D. Vlock, J. C. Lee, A. Kropec-Huebner, G. B. Pier, Pre-clinical and initial phase i evaluations of a fully human monoclonal antibody directed against the PNAG surface polysaccharide on *Staphylococcus aureus*. Abstracts of the 50th ICAAC 2010; Abstract G1-1654/329., (2010).

46. X. Yu, R. Prados-Rosales, E. R. Jenny-Avital, K. Sosa, A. Casadevall, J. M. Achkar, Comparative evaluation of profiles of antibodies to mycobacterial capsular polysaccharides in tuberculosis patients and controls stratified by HIV status. Clin. Vaccine Immunol. 19, 198–208 (2012).

47. L. Brown, J. M. Wolf, R. Prados-Rosales, A. Casadevall, Through the wall: extracellular vesicles in Gram-positive bacteria, mycobacteria and fungi. Nat. Rev. Microbiol. 13, 620–630 (2015).

48. J. Ludbrook, Multiple comparison procedures updated. Clin. Exp. Pharmacol. Physiol. 25, 1032–1037 (1998).

